# Revealing rhythm categorization in human brain activity

**DOI:** 10.1101/2024.12.04.626390

**Authors:** Francesca M. Barbero, Tomas Lenc, Nori Jacoby, Rainer Polak, Manuel Varlet, Sylvie Nozaradan

## Abstract

Humans across cultures show an outstanding capacity to perceive, learn, and produce musical rhythms. These skills rely on mapping the infinite space of possible rhythmic sensory inputs onto a finite set of internal rhythm categories. What are the brain processes underlying rhythm categorization? We used electroencephalography (EEG) to measure brain activity as human participants listened to a continuum of rhythmic sequences characterized by repeating patterns of two inter-onset intervals. Using frequency and representational similarity analyses, we show that brain activity does not merely track the temporal structure of rhythmic inputs, but, instead, automatically produces categorical representation of rhythms. Surprisingly, despite this automaticity, these rhythm categories do not arise in the earliest stages of the ascending auditory pathway, but show strong similarity between implicit neural and overt behavioral responses. Together, these results and methodological advances constitute a critical step towards understanding the biological roots and diversity of musical behaviors across cultures.

## INTRODUCTION

A fundamental function of the brain is to enable adaptive behavior in an environment full of remarkably diverse, dynamic sensory signals. Specifically, although constantly stimulated with a wide range of inputs, the brain does not treat each sensory input as a novel, unique event - a process that would be overwhelming for the organism – but instead categorizes it^1–5^. Perceptual categorization thus constitutes a core brain function, allowing to produce different responses to inputs belonging to different categories (*discrimination*) and, at the same time, identical responses to inputs belonging to the same category (*generalization*)^6^.

A striking illustration of this phenomenon is the central role categorization plays in human social interaction through musical rhythm^7–9^. In music, rhythm usually refers to patterns of durations between successive sensory events (i.e., inter-onset intervals between successive notes, or strokes of a percussion instrument). For example, we easily recognize and reproduce the rhythm of “Jingle Bells” or the stomp-stomp-clap groove of “We Will Rock You” as a prototypical rhythm composed of three time intervals, where the first and second intervals are simply twice shorter than the third interval (i.e., “Jin-” and “gle” vs. the twice longer “Bells”; same for “stomp” and “stomp” vs. the twice longer “clap”), hence the 1:1:2 ratio code often used to describe it (Suppl. Audio 1). Remarkably, we can easily recognize the rhythm of “Jingle Bells” even when the performed version deviates from a perfect template-like rendition of the time intervals. We are also able to discriminate this 1:1:2 rhythm from other distinctive three-interval rhythms, such as the so-called “tresillo” or “zouk” that is pervasive in many Afro-diasporic music genres over the world^10–13^ (with a prototypical 3:3:2 ratio; Suppl. Audio 2).

Hence, rhythm categorization enables us to recognize musical rhythms, by allowing an infinite space of possible rhythmic sensory inputs to be carved up into a finite set of internal categories. Critically, the existence of rhythm categories has been corroborated empirically using a number of behavioral paradigms including identification, discrimination, and reproduction^14–24^, and appears to be a phenomenon widely found across cultures^10^. Moreover, rhythm categories have been regarded as closely tied to the concept of a prototype. Rhythm prototypes are thought to represent specific patterns of relative durations, that is, specific rhythmic ratios in reference to which other rhythmic stimuli would be recognized, thus acting as a “perceptual magnet”^25^ or “attractor”^17^.

What are the biological processes underlying rhythm categorization? One view is that rhythm categories stem from hard-wired neurobiological predispositions constraining internal representations of rhythmic inputs. In particular, it has been proposed that rhythms corresponding to mathematically simple ratios (i.e., small integer ratios based on a grid of equal time intervals and their grouping in twos, such as the 1:1:2 rhythm of “Jingle Bells”^18^) should be universally privileged as a direct consequence of fundamental dynamics of neural assemblies^26^. This view has been recently corroborated by large-scale behavioral research showing small-integer ratio categories to be universally present across cultures^10^. This view also predicts small-integer ratio categories to emerge automatically at early stages of the sensory processing pathway.

However, a growing body of work points towards rhythm categorization as a plastic function, reflecting enculturation and social learning^22,27,28^. This view is supported by converging behavioral and modelling evidence that rhythm categories are not fully predetermined by hard-wired neurobiological processes^15,20^, but are shaped by culture-specific and individual experience^10,18,22,24,28,29^. Relatedly, rather than emerging spontaneously at early processing stages, rhythm categorization may be subserved by a plastic network of higher-level sensory, motor, and associative cortices, whose activity is shaped by the distribution of rhythms in the environment^22,27,28^. Moreover, this brain network could be selectively engaged depending on task demands^30^ (such as during sensorimotor reproduction or explicit discrimination; see ^4,31–33^).

Therefore, clarifying the interplay between hard-wired mechanisms and culture-driven neural plasticity appears a critical step to understand how socially meaningful categories of rhythm are produced and transmitted. Yet, this endeavor has proven particularly challenging so far, due to the lack of task-independent measures to capture rhythm categorization from neural responses. More broadly, task-free measures are ultimately key to probe rhythm categorization across the lifespan, cultures and species, and address long-standing questions regarding the nature and underlying mechanisms of human rhythmic behaviors. Here, we address this gap by providing, to our knowledge, first neural evidence for rhythm categorization and underlying rhythm prototypes. To this end, we developed a novel methodological approach combining (i) electrophysiology, (ii) frequency-domain analysis^34–37^ and (iii) the Representational Similarity Analysis framework (RSA)^38,39^, hereafter: *frequency-RSA* (fRSA).

Using this novel approach, we provide first direct evidence for neural categorization of rhythm in humans. Specifically, we show that brain activity captured with surface electroencephalography (EEG) goes beyond mere tracking of acoustic temporal features of the rhythmic inputs and, instead, exhibits categorical representations. Moreover, we show that these neural rhythm categories emerge automatically, without any related explicit task, yet they are remarkably similar to the categorical structure reflected in sensorimotor reproduction of the same stimuli. Surprisingly, despite this automaticity, these rhythm categories do not arise in the earliest stages of the ascending auditory pathway, as revealed by biomimetic modeling. Therefore, by ruling out that the process of rhythm categorization merely reflects motor, instructional or decisional biases, our results advance a critical step forward in the understanding the nature and neural pathways underlying this function fundamental to the human experience of music.

## RESULTS

Using scalp electroencephalography (EEG), we recorded brain activity of healthy adult participants (N = 18) as they listened to different sequences, each made of a repeated rhythmic pattern (750-ms pattern duration), while carrying out an unrelated volume-change detection task. Each pattern consisted of two identical tones, which, when looped, yielded a repeating sequence of two time intervals. Across conditions, we manipulated the ratio between the duration of the first and the second time interval to span the continuum of two-interval rhythms ranging from isochrony (1:1 interval ratio) to patterns of long-short intervals with 2:1 ratio (hereafter also expressed as ratios from 0.50 to 0.67, respectively, when dividing the first interval by the sum of the two intervals; Fig. 1; Suppl. Table 1; Suppl. Audio 3).

**Figure 1.**
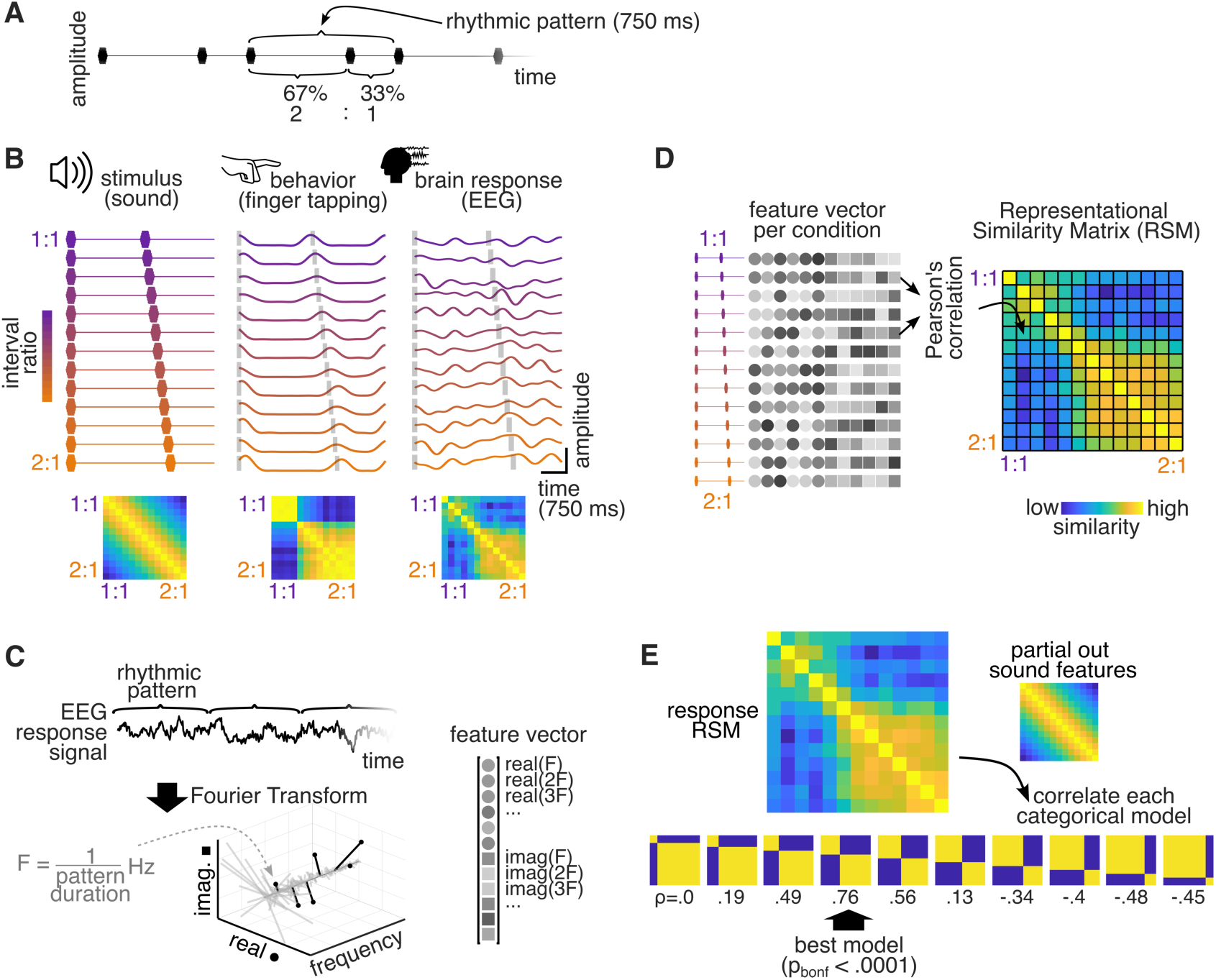
Overview of the experimental design, the frequency-RSA approach, and rhythm categories as obtained in an example participant. A. Example excerpt depicting the time course of the auditory stimulus from one of the conditions, showing several repetitions of the 750-ms long two-interval pattern used to construct the 22.5-s long stimulus sequences. Individual tones are indicated with black markers. One repetition of the rhythmic pattern is highlighted using brackets above the stimulus. The two constituent inter-onset intervals are indicated with brackets below the stimulus, showing the percentage of the total pattern duration spanned by each interval, as well as the ratio of the two intervals (here 2:1). B. Stimulus design and responses of an example participant. Left: Time course of the 750-ms two-interval pattern used to construct the stimulus sequence in each condition. Colored markers indicate positions of the tones making up the pattern. The ratio of the two constituent intervals gradually changes across the 13 conditions, from 1:1 (equal intervals) to 2:1 (one interval twice longer than the other), as indicated by the color gradient. Below is the acoustic similarity between pairs of stimuli depicted in the form of a Representational Similarity Matrix (RSM). The matrix structure reflects the smooth decrease in similarity across the interval ratios of the conditions. Middle: Average time course of the tapping force generated by an example participant when synchronizing the tapping with the rhythmic stimuli in each condition. For each condition, tone onsets are indicated by vertical grey bars. The tap-force RSM below was built using the frequency Representational Similarity Analysis (fRSA) method. The tap-force RSM shows that the tapping responses are mutually similar when the stimulus is near 1:1 or 2:1 ratio, with a sharp dissimilarity boundary separating the two categories. Right: Trial-averaged EEG responses of an example individual participant recorded while listening to the same stimuli without overt movement. The neural RSM below shows a categorical structure that is highly similar to the tapping response. C. Feature extraction. The top panel shows a segment of an example EEG signal in a representative condition depicting neural response to several repetitions of the rhythmic pattern (indicated by brackets). The complex-valued spectrum of the EEG obtained with Fourier Transform is shown below. The response is concentrated in a-priori known frequency bins corresponding to 1 / rhythmic pattern duration and harmonics (highlighted in black) and can be isolated from noise broadly distributed across frequencies. Real and imaginary coefficients at response frequencies are concatenated into a feature vector shown on the right. The magnitudes of real and imaginary coefficients (depicted with circles and squares respectively) are indicated using gray scale. D. Representational similarity matrix (RSM) built by correlating feature vectors across all pairs of conditions. E. Fitting categorical models to the response data. The neural RSM of an example individual participant is correlated with each candidate categorical model RSM (differing in the location of the category boundary) after removing the contribution of the acoustic model RSM. The p-value is obtained using permutation testing. Icon sources: “sound” by PureSolution, “EEG” by Aenne Brielmann, and “finger” by iconcheese from the Noun Project under CC BY 3.0 license.

We used this rhythm continuum based on previous results consistently showing that rhythms from this continuum are perceived by Western participants as two discrete categories separated by a sharp perceptual boundary^14,15,22^. In addition, each of these categories has been characterized by a prototypical rhythmic pattern, one centered on 1:1 and the other close to (but not necessarily at) 2:1 ratio^16–19^. There is abundant evidence that sensorimotor synchronization and reproduction of rhythms falling in each of these categories are distorted towards the corresponding prototype, thus broadly compatible with the concept of a perceptual attractor from dynamical systems literature^16,17^ as well as a Bayesian concept of a perceptual prior^22,27^.

To obtain a behavioral measure of rhythm categorization, we also asked the same participants to tap their finger on a response sensor in the best possible synchrony with the stimuli (in blocks separate from the EEG recordings).

### Behavioral evidence for rhythm categorization

#### Inter-tap interval ratio analysis

First, we tested whether rhythm categorization was evident in the behavior by analyzing the inter-tap intervals (ITI) produced in each condition when participants were instructed to tap the finger in synchrony with the rhythmic inputs (Fig. 2A). To this aim, we calculated the ratio of the first over the sum of the first and second ITI separately for each of the 90 (30 per sequence x 3 trials) repetitions of the 750-ms long rhythmic pattern. After cleaning (see Methods), ITI ratios were averaged across rhythm repetitions separately for each condition and participant (mean number of averaged ITI ratios per condition = 64, range = 16-84).

**Figure 2.**
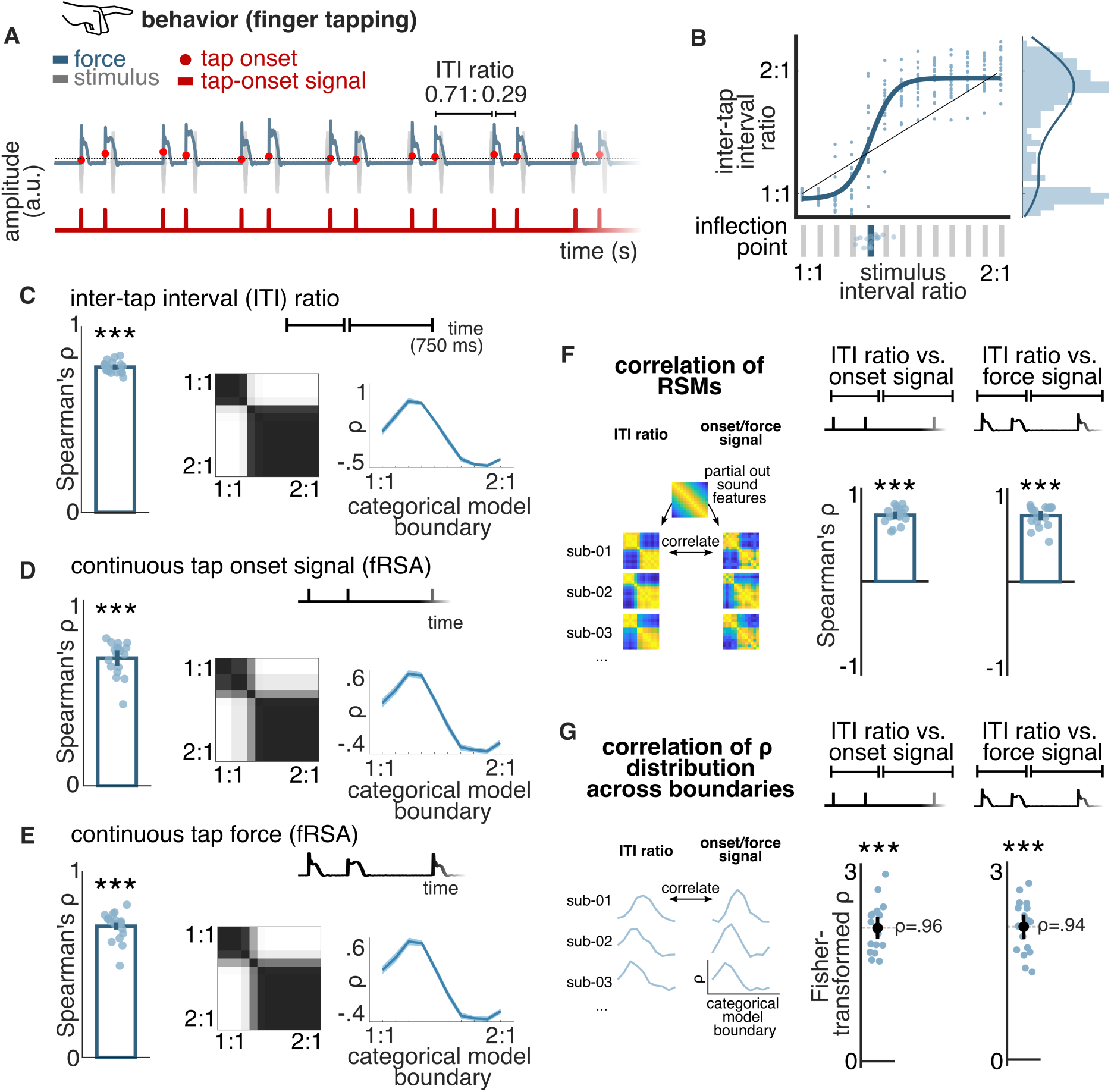
Analyses of behavioral responses show consistent categorization across participants and tapping signals. A. Example segment of a continuous tapping force signal (in blue). Tap-onset time points are indicated with red circles superimposed on the continuous tapping force. Two inter-tap intervals (ITIs) and their ratio within an example repetition of the rhythmic pattern are highlighted with brackets. The corresponding continuous tap-onset signal is shown in red below. B. Average ITI ratio as a function of stimulus inter-onset interval (IOI) ratio. Responses from individual participants are shown as blue circles. The black diagonal line corresponds to identical ITI and IOI ratios, i.e., undistorted reproduction. The blue line depicts a sigmoid function fit to data pooled across all participants, and its 50% threshold location is shown as a vertical blue line in the panel below. The same parameter estimated separately for each participant is plotted using blue circles. The pooled distribution of ITI ratios is shown as a histogram on the right and the estimated density is indicated with a blue line. C. The left panel shows Spearman’s correlation of the ITI RSM (as built from absolute differences between average ITI ratios produced in each condition) with the best-fitting categorical model RSM, obtained separately for each participant. Blue circles indicate data from individual participants and error bars represent a 95% confidence interval. Asterisks indicate a significant permutation test at the group level (Bonferroni corrected, *p < 0.05, **p < 0.01, ***p < 0.001). The middle panel shows an overlay of best-fitting categorical model RSMs across individual participants, highlighting the remarkable consistency of the estimated category boundary. Distribution of correlations across all possible categorical models differing in the location of the category boundary is shown on the right (averaged across participants, shaded regions indicate 95% confidence interval). D. Same as panel C, but for the continuous tap-onset signal (considering harmonics up to 16 Hz). E. Same as panel C, but for the continuous force signal (considering harmonics up to 16 Hz). F. Participant-wise correlation of the behavioral RSM based on ITI ratios and the corresponding RSM based on continuous tap-onset signals (left) and continuous tap-force signals (right). The shared information driven by stimulus features was removed by partialling out the acoustic model RSM. Data from individual participants are shown as blue circles (filled circle indicates a significant permutation test at the individual participant level). Error bars represent a 95% confidence interval. Asterisks indicate a significant permutation test at the group level (*p < 0.05, **p < 0.01, ***p < 0.001). G. Fisher-transformed correlation between the distribution of correlations across all possible categorical models obtained for the ITI RSM and the tap-onset RSM (left) or the tap-force RSM (right). Blue circles correspond to individual participants, error bars represent a 95% confidence interval, and asterisks indicate a significant t-test against zero (*p < 0.05, **p < 0.01, ***p < 0.001). Icon sources: “finger” by iconcheese from the Noun Project under CC BY 3.0 license.

In line with previous studies^17,19,21,22^, the produced ITI ratios did not faithfully reproduce the ratios of the corresponding stimuli (Fig. 2B). Instead, they revealed two categories, whereby the tapping near the 1:1 and 2:1 conditions (which can also be expressed as 0.50 and 0.67) exhibited a ratio close to 1:1 and 2:1 respectively. Indeed, the distribution of ITI ratios across conditions was significantly better explained by a sigmoidal compared to a linear model (p = 0.002, Wilcoxon rank sum test comparing leave-one-participant-out cross-validated R^2^). Moreover, the mean inflection point of the sigmoidal fit across participants did not overlap with the mathematical mid-point of the ratio continuum tested in the current study (i.e., 0.58, but corresponded instead to a median ratio of 0.56 (bootstrapped 95% CI = [0.56, 0.56]), that is, relatively closer to the 1:1 ratio edge of the continuum, thus revealing a smaller and a bigger category^14^. The two categories were also apparent in the distribution of produced ITI ratios collapsed across all conditions and participants that was significantly different from a uniform distribution (χ^2^(12) = 113.51, p < 0.0001). Taken together, the obtained results corroborated a well-established effect in the literature^18,40,41^.

#### RSA of inter-tap interval ratios

The categorical representation evident in the produced ITI ratios was further corroborated using the Representational Similarity Analysis (RSA) framework. We extracted individual representational geometries directly from the ITIs: for each participant, we built a Representational Similarity Matrix (RSM) of the ITIs based on pairwise absolute differences in the average interval ratios produced across conditions. The obtained ITI RSMs were then correlated with theoretical models of categorization, while controlling for the variance explained by acoustic features (Fig. 1E; see Methods).

As shown in Figure 2C, the ITI RSMs significantly correlated with the categorical models at the group level (permutation test, *ρ* = 0.78, p_bonf_ = 0.001) and at the individual level (when evaluated for each participant separately) for all participants (permutation test, ps_bonf_ < 0.05; see also Fig. S1). The best-fitting categorical models were consistent across individuals, and the observed category boundary (median ratio 0.55, bootstrapped 95% CI = [0.55, 0.56]) was compatible with the one identified with ITI ratio analysis.

#### Frequency-RSA of tapping onset time series

Next, we extended the ITI-based findings by testing whether a similar categorical structure could be observed when representing the series of tap onsets as a continuous signal. This constitutes a pivotal shift from the analysis of time intervals defined by discrete temporal markers to the analysis of continuous data where the identification of temporal markers is less straightforward (e.g., as in surface EEG). To this end, we applied the frequency-RSA (fRSA) approach to the tapping responses represented as continuous time-varying signals with a unit impulse at the onset time of each executed tap (see Fig. 2A). Notably, and in contrast with ITI ratios, such representation is sensitive to the stability of the produced intervals over trials, thus constituting a more comprehensive measure to capture categorization in continuous signals.

For each participant and condition, the spectrum of trial-averaged continuous tap-onset signals was obtained using Fourier Transform. A critical advantage of using frequency-domain analysis in the current experimental design is that it allows for isolating responses to the repeated rhythmic pattern from the recorded continuous signals with high objectivity and signal-to-noise ratio^34–37^ (Fig. 1C). Indeed, in line with well-established principles of frequency-tagging^34,42,43^, the set of complex Fourier coefficients at the frequency of the periodically presented rhythmic pattern and harmonics effectively describes the properties of *any* response that *consistently repeats* across pattern repetitions.

As expected, based on this principle, the obtained spectra averaged across all participants and conditions exhibited peaks of magnitude at frequencies corresponding to the repetition rate of the rhythmic pattern and harmonics (i.e., *f* = 1/0.75 s = 1.33 Hz, 2*f* = 2.66 Hz, and so on; Fig. S2). The magnitude of the response at these a priori determined frequencies was significantly above the local noise baseline^44–46^ for harmonics up to 16 Hz (z > 3.09, i.e., p < 0.001, one-tailed, signal > noise), which were thus further considered as frequencies of interest (Fig. S3). Responses at these frequencies of interest were used to build an RSM for each participant. First, the real and imaginary Fourier coefficients obtained at harmonics up to 16 Hz were concatenated to form a feature vector for each condition (Fig. 1C), thus preserving maximal amount of information (including amplitude and timing information) relevant to the responses. Subsequently, an RSM was built to reflect the similarity of feature vectors across conditions (Fig. 1D).

There was a significant correlation between tap-onset RSMs and categorical models at the group level (permutation test, *ρ* = 0.69, p_bonf_ = 0.001), as well as at the individual level in all participants (permutation test, ps_bonf_ < 0.05; Fig. 2D; see also Fig. S1). As for the results obtained with the RSA applied to ITI ratios, there was a marked cross-participant consistency in the category boundary that best explained the RSMs obtained with the novel fRSA applied to the continuous tap-onset signals (Fig. 2D, median boundary at ratio 0.56, bootstrapped 95% CI = [0.55, 0.56]).

The correlation between the RSM and the best-fitting categorical model selected for each participant was significantly smaller when considering the tap-onset signals rather than ITIs (paired t-test, p < 0.001, BF_10_ = 76). This was not surprising given that frequency-domain analysis of tap onset signals is more conservative as compared to ITI-ratio analysis which comprised: (i) a cleaning step whereby responses to pattern repetitions where the participant was not complying with the synchronization task were discarded from further analyses, and (ii) an inherent time-warping of the response to each pattern repetition that corrects for tempo drifts (as the absolute durations of the two ITIs produced on each pattern repetition are normalized by taking a ratio). Nonetheless, there was a remarkable similarity between the RSMs obtained from ITIs and from the tap-onset signals (Fig. 2F, group-level permutation test, *ρ* = 0.72, p = 0.0001). Moreover, the relative distribution of correlation coefficients across theoretical models with different locations of the category boundary was remarkably similar between the two analyses (Fig. 2G, mean *ρ* across participants = 0.96, one-sample t-test against zero, p < 0.0001). These findings demonstrate that fRSA can reveal categorization encoded in the temporal structure of a response without relying on the extraction of discrete temporal markers.

#### Frequency-RSA of continuous tapping force

To provide further evidence for rhythm categorization as captured with continuous signals, we applied the fRSA analysis to the continuous force signals obtained directly from the tapping sensor. In addition to the temporal arrangement of the taps, the force signal also captures their relative accentuation, thus potentially offering yet another dimension that could reflect categorization of rhythmic inputs. Inspecting the grand-average magnitude spectrum of the tapping force revealed peaks at the rate of the rhythmic pattern repetition up to 16 Hz (z > 3.09, i.e., p < 0.001, one-tailed, signal > noise; Fig. S3; see also Fig. S2). We thus used the real and imaginary Fourier coefficients at these frequencies of interest to build a tap-force RSM for each participant.

At the group level, the obtained tap-force RSMs showed clear correspondence with a categorical representational structure (permutation test, *ρ* = 0.71, p_bonf_ = 0.0001), which was also significant in each individual participant (permutation test, ps_bonf_ < 0.05; Fig. 2E, see also Fig. S1 and Fig. 3B), and consistent across participants (median boundary at ratio 0.55, bootstrapped 95% CI = [0.55, 0.56]). The average correlation with a categorical model was significantly smaller for RSMs built by applying fRSA to the force signal, as compared to the RSMs obtained from ITI ratios (paired t-test, p < 0.001, BF_10_ = 70). Yet, similarly to the results obtained with fRSA of tap-onset signals reported above, the categorical structure captured by the tap-force fRSA was strikingly consistent with the one identified using ITI ratios. Specifically, the corresponding RSMs obtained by each method highly correlated within participants (Fig. 2F, group-level permutation test, *ρ* = 0.72, p = 0.0001), and revealed a similar position of the category boundary (Fig. 2G, mean *ρ* across participants = 0.94, one-sample t-test against zero, p < 0.0001).

**Figure 3.**
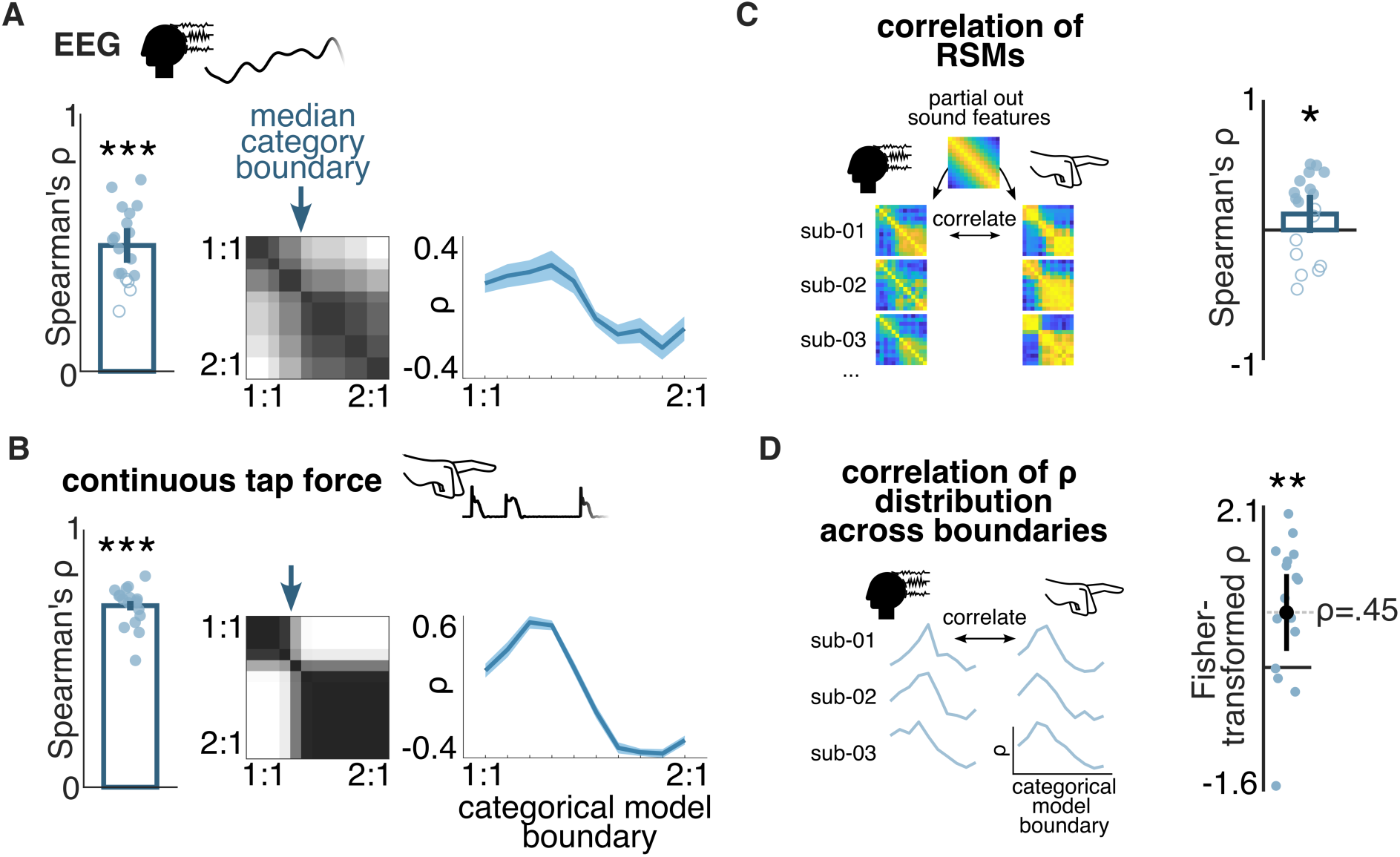
Convergent evidence for rhythm categorization from neural and behavioral responses, as captured with fRSA. A. fRSA analysis of EEG responses (responses from 64 channels, considering frequencies up to 8 Hz). Left: blue circles show Spearman’s correlation of the EEG response RSMs with the best-fitting categorical model, obtained separately for each participant. Filled circles indicate a significant permutation test at the individual participant level, and asterisks indicate a significant permutation test at the group level (Bonferroni corrected, *p < 0.05, **p < 0.01, ***p < 0.001). Middle: overlay of best-fitting significant categorical models across participants, with median category boundary indicated with a blue arrow. Right: distribution of correlations across all possible categorical models differing in the location of the category boundary (averaged across participants, shaded regions indicate 95% confidence interval). B. Same as panel A but applied to continuous tapping force (considering frequencies up to 16 Hz). C. Participant-wise correlation of the neural RSM and the corresponding tap-force RSM. The shared information driven by stimulus features was removed by partialling out the acoustic model RSM. Data from individual participants are shown as blue circles (filled circle indicates a significant permutation test at the individual participant level). Error bars represent a 95% confidence interval. Asterisks indicate a significant permutation test at the group level (*p < 0.05, **p < 0.01, ***p < 0.001). D. Fisher-transformed correlation between the distribution of correlations across all possible categorical models obtained for the neural RSM and the tap-force RSM. Blue circles correspond to individual participants, error bars represent a 95% confidence interval, and asterisks indicate a significant t-test against zero (*p < 0.05, **p < 0.01, ***p < 0.001). Icon sources: “EEG” by Aenne Brielmann, and “finger” by iconcheese from the Noun Project under CC BY 3.0 license.

These results thus highlight the fRSA approach as a conservative yet sensitive method to capture categorical structures from continuous smooth signals beyond relying on discrete homogenous temporal markers. Notably, these results also indicate that including additional information about the way each tap movement is executed (here, including both tap onset and force information) yields representational structures overall consistent with those obtained with tap timing alone. In line with prior work^16^, these results provide further support to the view that the timing of motor responses elicited during a sensory-motor synchronization task is the key dimension that reflects rhythm categorization in human adults.

#### Identifying the underlying prototypes

Using fRSA, our results provided evidence for categorical structure in continuous tapping signals. From this, we then aimed at moving a step forward and informing on the actual structure of the responses making up each identified category. Indeed, the categorical structure observed here could have been driven by any response feature of the tapping signal that systematically differed across the two categories (i.e. discriminated) and remained consistent for conditions within each category (i.e. generalized). In principle, the relevant feature could have been a particular pattern of inter-tap intervals, and, in case of the continuous force signal, also a particular pattern of accentuation.

To capture the structure of the response making up each identified category, we assessed for each condition the similarity between the response and a set of prototypical signals. The prototypes consisted of 76 two-interval pattern templates (with unit impulse at the onset time; 750 ms pattern duration, as in the stimuli) whose ratios were equally spaced between 0.50, i.e., the 1:1 edge of the condition continuum tested here, to 0.84, i.e., beyond the 2:1 ratio (0.67) corresponding to the other edge of the condition continuum. This set of prototypes thus corresponded to a wide and fine gradient of two-interval ratios (note that ratios from 0.84 up to 0.99 were not included in this set of prototypes as these ratios would result in shortest interval durations shorter than 127.5-ms, thus likely reaching motor constraints for one-to-one sensorimotor synchronization of finger tapping in non-musicians^47^).

The Fourier spectrum of each template signal revealed a characteristic distribution of magnitudes across the frequencies of interest corresponding to the rate of rhythmic pattern repetition and harmonics (Fig. 4A). Unlike complex Fourier coefficients, the distributions of magnitudes reflect the temporal structure of each individual prototype signal characterized by a repetition of a consistent trajectory separated in time by the given interval ratio, yet without additional assumptions about the particular shape or phase lag of the repeated trajectory (see also^48^). In other words, these prototypical distributions of magnitudes across frequencies of interest can be considered a set of hypotheses about the temporal structure of a/symmetries in a signal.

**Figure 4.**
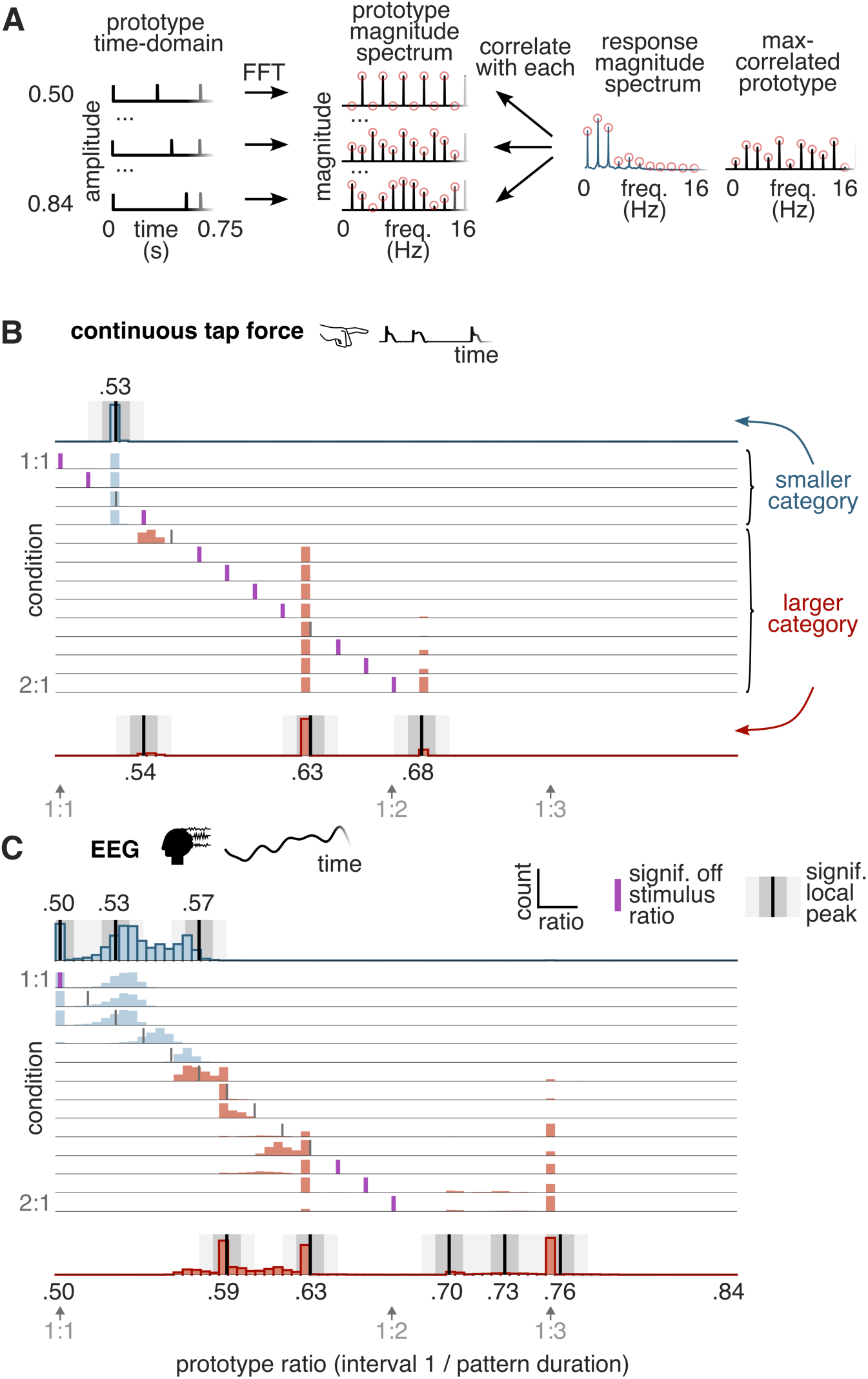
Analysis of similarity with two-interval prototypes: identified prototypes deviate from the mathematically simplest ratios, in both neural and behavioral responses. A. Schematic of the analysis to evaluate similarity between the response and a set of prototypical template signals. The time-domain representation of several example prototypes is shown on the left. The prototypes were made of two repeated impulses, arranged to create a repeating 750-ms long pattern of two intervals with ratios spanning 0.50 - 0.84. The magnitude spectra of the example prototypes are shown in the middle, with harmonics of the repeating pattern rate highlighted with red circles. A grand average response magnitude spectrum from an example condition (obtained in one iteration of the bootstrapping procedure) is shown on the right. The vector of magnitudes at frequencies of interest (red circles) is correlated with the corresponding vector of each prototype, and the maximally correlated prototype is selected. B. Bootstrapped distribution of maximally correlated prototypes for the continuous tap-force responses (considering frequencies up to 16 Hz). A histogram of best-correlating prototypes (x-axis) is shown separately for each condition (y-axis). Conditions corresponding to the smaller and bigger category identified in the fRSA analysis are shown in blue and red, respectively. The vertical line in each row marks the stimulus ratio in the corresponding condition. When magenta, this line indicates a condition with significantly greater distribution density outside vs. inside a narrow window centered on the stimulus ratio. A distribution collapsed across the conditions within each category is shown on the top and bottom, respectively. Positions of significant local density peaks are depicted with vertical black lines, flanked by grey rectangles indicating snippets of the histogram that were used to quantify prominence of the particular peak (calculated as density inside the darker rectangle minus the density inside the lighter rectangle). Grey arrows below the plot indicate positions of small integer ratios on the x-axis. C. Same as panel B, but for EEG responses (based on a pool of 9 fronto-central channels, and frequencies up to 8 Hz; note that the median category boundary estimated for the EEG responses is shifted one condition toward 1:1, as compared to the tapping responses). Icon sources: “EEG” by Aenne Brielmann, and “finger” by iconcheese from the Noun Project under CC BY 3.0 license.

To test which prototype offered the best characterization of the tapping response in each condition, we correlated the vector obtained from the magnitude spectrum of each prototype with the same vector obtained from the magnitude spectrum of the continuous tap-force signal averaged across all participants, separately for each condition. Grand average was used instead of individual participants’ spectra to further improve signal-to-noise ratio in view of optimizing identification of potential underlying prototypes. For each condition, we obtained a distribution of the maximally correlated prototype by bootstrapping participants taken for the calculation of the grand-average magnitude spectrum.

Importantly, the identified prototypes were not gradually changing across conditions, as would be expected if the response was following the temporal structure of the corresponding auditory stimuli (Fig. 4B). Indeed, the distribution of best-correlated prototypes was significantly concentrated away from the stimulus ratio in 11 out of the 13 conditions (ps_bonf_ < 0.05, bootstrap test).

We then assessed whether responses within each of the categories, as identified by the fRSA approach detailed above, showed high similarity to particular prototypes. To this aim, we created marginal distributions by collapsing the distribution of maximally correlated prototypes across conditions within the smaller and bigger category, respectively. Local peaks in these marginal distributions were identified by sliding a narrow window through the prototype continuum and using a bootstrap analysis to quantify whether the distribution inside the window was higher than in its local neighborhood^49,50^ (see Methods; Fig. 4).

On the one hand, in the four conditions corresponding to the smaller category (i.e., between stimulus ratios 0.50 and 0.54), the distribution of prototypes maximally correlated with the tapping response peaked at a ratio near 0.53 (p_bonf_ = 0.03). On the other hand, for the eight conditions corresponding to the bigger category (i.e., between 0.56 and 0.67), the tapping seemed to be best characterized by a prototype with ratio near 0.63 (p_bonf_ = 0.03), with smaller but significant peaks appearing near 0.68 (p_bonf_ = 0.03) and 0.54 (p_bonf_ = 0.03, yet limited to a single condition).

Importantly, a qualitatively identical profile of results emerged when the same analysis was run on the continuous tap-onset signals, confirming robustness of the analysis to the particular shape of the repeated signal trajectory. Together, these observations point toward underlying prototypes that would not align with small, mathematically simple, integer ratios (i.e., 0.50 and 0.67, corresponding to 1:1 and 2:1, respectively). Instead, the participants seemed to soften/sharpen the produced interval ratio in a way broadly consistent with observations from other behavioral studies^16–18^.

#### Summary of behavioral results

All in all, the results of our tapping analyses (i) confirm that the set of two-interval rhythms used in the current study elicited internal representation of two separable rhythm categories, (ii) validate the fRSA method as a robust and sensitive tool to identify rhythm categories using continuous time-varying response signals beyond the need to extract discrete temporal markers, and (iii) point toward underlying rhythm prototypes that deviate the mathematically simplest ratios, in line with previous studies^16–18^.

### Neural evidence for rhythm categorization

#### Frequency-RSA of EEG responses

We recorded brain activity using electroencephalography (EEG) as participants listened to the rhythmic stimuli without performing any overt movement. As for the tapping, trial-averaged preprocessed EEG responses were transformed into the frequency domain using Fourier Transform. Here, we only considered frequencies of interest up to 8 Hz, as this frequency cutoff captured all significant responses at consecutive harmonics of the rhythm repetition rate as observed in the grand-average EEG magnitude spectrum (i.e., first 6 harmonics of the rhythmic pattern repetition; z > 3.09, i.e., p < 0.001, one-tailed, signal > noise; Fig. S3, see also Fig. S2). Higher frequencies were also excluded to avoid distortion of the EEG spectra by the alpha activity artifact (approximately between 8 and 12 Hz^51–53^).

For each participant, we built a neural RSM based on the similarity of real and imaginary Fourier coefficients at the frequencies of interest concatenated across all 64 EEG channels, thus accounting for any individual differences in response topography. The neural RSMs exhibited significant categorical structure at the group level (permutation test, *ρ* = 0.49, p_bonf_ = 0.001; Fig. 3A). Moreover, we were able to observe significant correlation with a categorical model at the individual level in most participants (14 out of 18; permutation test, ps_bonf_ < 0.05; Fig. 3A, see also Fig. S1 and Table S2).

Notably, we observed comparable evidence of neural categorization when limiting the analysis to responses averaged across 9 fronto-central channels (Fig. S4). This pool of channels was selected based on the fact that they have been shown to consistently capture EEG responses to repeating acoustic rhythms across previous studies^36,54–56^. Analogously, these channels also showed the highest overall response magnitude averaged across all participants and conditions in the current study (Fig. S3). This result suggests that accounting for individual differences in response topography is not critical to reveal categorical encoding of auditory rhythmic inputs. Instead, it appears that spatiotemporal dynamics captured in the time course of brain activity at a single scalp location (with high signal-to-noise ratio) are sufficient to efficiently capture the representational geometry relevant for rhythm categorization.

Finally, we also observed comparable evidence of neural categorization when neural RSMs were built using the time course of EEG responses over the duration of the rhythmic pattern after bandpass filtering and averaging these responses across all repetitions of the pattern. Indeed, these convergent results are expected, as such time domain averaging acts as a comb filter that effectively isolates frequencies corresponding to pattern repetition and harmonics, which is a fundamental rationale underlying fRSA (Fig. S5).

Taken together, these results provide evidence for neural categorization of rhythm, whereby neural activity elicited without related temporal task or overt movement represents the acoustic continuum of two-interval rhythms as two distinct categories.

#### Frequency-RSA of auditory nerve model responses

In our fRSA analysis of the EEG responses reported above, we controlled for potential contribution of low-level tracking of the sensory input to the obtained categorical structure by partialling out the acoustic RSM (Fig. 1E). However, it remains unclear whether the observed transformation from acoustic features towards a categorical representation of rhythm reflects a higher-level, possibly cortical neural processing, and to what extent it could be shaped by lower-level nonlinearities of subcortical neurons from the ascending auditory pathway. We explored this possibility by taking advantage of a well-established biologically plausible model of the auditory periphery to simulate responses elicited by our acoustic stimuli in a set of auditory nerve fibers^57^.

The model provided faithful simulation of physiological processes associated with cochlear nonlinearities, inner hair cell transduction process, the synapse between the hair cell and the auditory nerve, and the associated firing rate adaptation. For each condition, we simulated the time course of instantaneous firing rate in the auditory nerve elicited by the corresponding rhythmic sequence (Fig. S6A). An auditory-nerve RSM was then obtained from these simulated responses by applying the same fRSA analysis as for the EEG responses above.

Crucially, the auditory-nerve RSM closely resembled the acoustic RSM (Fig. S6B), indicating that early subcortical representations mainly follow the temporal structure of the stimulus sequences. More specifically, the obtained auditory nerve RSM did not show significant rhythm categorization (no significant correlation with a categorical model; permutation test, *ρ* = 0.33, p_bonf_ = 0.18), in contrast with the EEG responses. Most importantly, when accounting for structure explained by the auditory-nerve RSM using partial correlation, the EEG responses still showed categorical representation of rhythm significant both at the group level (permutation test, *ρ* = 0.51, p_bonf_ = 0.001) and at the individual level in 16 out of the 18 participants (permutation test, ps_bonf_ < 0.05).

Together, these results suggest that neural categorization of rhythm does not likely arise as a product of nonlinear transformations occurring at the earliest, peripheral stage of the ascending auditory pathway.

#### Identifying the underlying neural prototypes

Using fRSA, we were able to observe categorical representation of rhythm in neural responses measured with scalp EEG. Based on this evidence, and similar to the behavioral analysis, we then moved a step forward by characterizing rhythm prototypes underlying the observed categories. To do so, we compared the neural response in each condition to the same set of prototypical ratio templates as used in the analysis of continuous tapping responses, i.e., a range of 76 prototypes corresponding to interval ratios between 0.50 and 0.84 (Fig. 4A). We considered magnitudes at the first 6 harmonics of the pattern repetition rate (i.e., up to 8 Hz), extracted from the grand-average magnitude spectrum of EEG activity in a pool of 9 fronto-central channels, to optimize signal-to-noise ratio. The obtained magnitude vectors were correlated with the set of prototypes, and a distribution of the best-correlated prototype in each condition was calculated by bootstrapping.

The observed distribution appeared to partly follow the interval ratios of the rhythmic patterns presented in each condition (Fig. 4C). This is not surprising, as the measured EEG response is expected to reflect a mixture of stimulus features and categorization, which are less efficiently disentangled here as compared to the fRSA analysis where stimulus features are partialled out. Crucially, the EEG response appeared to go beyond simple replication of the stimulus interval ratios, as suggested by significantly higher density of maximally correlated prototypes concentrated off rather than on the stimulus ratio in 4 out of 13 conditions (ps_bonf_ < 0.05, bootstrap test). Instead, high similarity tended to cluster near a small set of prototypes, that were evident after collapsing the distribution of best-correlated prototypes across conditions separately for each category identified with fRSA.

On the one hand, there seemed to be prominent similarity with prototypes corresponding to small integer ratios, indicated by significant peaks near 0.50 (i.e. isochrony; p_bonf_ = 0.03) or 0.75 (i.e. one interval three times longer than the other interval; p_bonf_ = 0.03). On the other hand, the structure of the EEG responses also appeared compatible with more complex interval ratios, including, for example, 0.53 (p_bonf_ = 0.03) and 0.63 (p_bonf_ = 0.03). Moreover, the neural categorical structure appeared to be somewhat more differentiated than the behavioral one. First, a prototype emerged at 0.59 (approximately corresponding to 4:3; p_bonf_ = 0.03). Second, the bias towards the prototype at 0.75, which lies outside the range of the stimuli as presented here (0.50–0.67 [1:1–1:2]) was not observed in the tapping. This prototype could reflect resonant frequencies emerging from fundamental oscillatory dynamics of neural assemblies^26^. These observations thus lend support to the view that neural activity elicited by rhythmic inputs may reflect a combination of acoustic features and internal biases that shape the representation towards a small set of prototypical rhythmic templates that do not systematically correspond to the mathematically simplest ratios.

### Convergence across behavioral and neural categorization of rhythm

Our results indicate that both behavioral and neural responses to the rhythmic inputs do not simply reflect acoustic features, but instead exhibit representational geometries consistent with the existence of two distinct rhythm categories, a smaller one spanning ratios from about 0.50 to 0.55 (1:1 - 1.2:1) and another, bigger, one spanning from 0.56 to 0.67 (1.3:1 – 2:1). However, is there a correspondence between the categories reflected in the brain and behavior?

To answer this question, we first assessed the overlap between the representations measured from the brain and behavior of each participant, by correlating the neural RSM (considering frequencies up to 8 Hz and all 64 channels) with the corresponding tap-force RSM (considering frequencies up to 16 Hz), while partialling out the shared similarity structure driven by the acoustic stimulus. The neural RSMs significantly correlated with the tap-force RSMs at the group level (permutation test, *ρ* = 0.12, p = 0.03), and this correlation was significant in 10 out of 18 participants (Fig. 3C). Similar results were obtained when correlating neural RSMs with tap-onset RSMs (group-level permutation test, *ρ* = 0.16, p = 0.007) or tap-ITI RSMs (group-level permutation test, *ρ* = 0.21, p = 0.003). These results suggest that the neural and behavioral responses share similar representational geometries beyond faithful encoding of acoustic features.

Next, we examined the location of the categorical boundary separating the two rhythm categories observed in the EEG and tap-force RSMs. Across participants, the location of this boundary in the best-fitting categorical model was remarkably similar for the EEG (median boundary at ratio 0.56, bootstrapped 95% CI = [0.54, 0.57]; Fig. 3A) and the tap-force responses (median boundary at ratio 0.55, bootstrapped 95% CI = [0.55, 0.56]; Fig. 3B).

This result was further corroborated by considering all theoretical models of categorization differing in the position of the category boundary, rather than a single best-fitting model. Indeed, the distribution of correlation coefficients obtained across all categorical models separately for each participant showed marked similarity between the neural and behavioral (tap-force) responses (mean *ρ* across participants = 0.45, one-sample t-test against zero, p = 0.01, Fig. 3D). Together, these observations highlight the consistency across neural and behavioral representations of the two-interval rhythms tested in the current study.

## DISCUSSION

The current study provides direct evidence for neural categorization of rhythm in humans. Specifically, we show that brain responses to rhythmic patterns do not merely reflect the physical temporal structure of the acoustic input. Rather, the structure of the neural responses across conditions is compatible with the existence of two distinct rhythm categories, consistent with behavioral measures from the same participants, and in line with a large body of prior behavioral work^14–23^. Moreover, the fact that neural categorization of rhythm emerges even in the absence of an explicit behavioral timing task suggests a largely automatic process.

### Probing rhythm categorization and its automaticity beyond behavioral measures

Thus far, research on rhythm categorization has been restricted to behavioral measures, due to the lack of a method allowing to capture rhythm categorization from neural data. Our results extend previous findings from discrimination and sensorimotor synchronization studies^10,14,22^, by showing that the brain automatically maps the continuous space of two-interval rhythms onto discrete rhythm categories also in the absence of a motor or timing-related task.

Crucially, this observation adds to the broad evidence arguing against the view that rhythm categorization arises in motion production from movement kinematics rather than from perceptual representations^58^. Indeed, prior work has revealed comparable rhythm categorization in sensorimotor tasks involving various modes of production (unimanual^19,21^ and bimanual tapping^59^, verbal reproduction^22^). Yet, beyond movement kinematic properties specific to each effector, different modes of movement executions could still, in principle, reflect constraints of higher-level, central motor control rather than perceptual representations^60^. However, without denying a role of movement for rhythm perception in general, a critical contribution of central motor control to rhythm categorization seems unlikely given robust evidence for rhythm categorization in perceptual discrimination tasks^14,20,22^.

Most importantly, we move a critical step beyond behavioral studies by showing here that profiles of discrimination/generalization compatible with perceptual categorization can emerge from neural activity even without engagement in related explicit judgement tasks that may be sensitive to decisional and cognitive factors potentially driven by task demands^4,33^. Our results thus reinforce the view of rhythm categorization as a largely automatic function of the human brain. The notion that this function might not be under volitional control is also compatible with previous observations whereby even highly trained musicians under strict instructions cannot avoid categorical distortions in sensorimotor reproduction of rhythm^16–18,41^.

### Candidate neural mechanisms underlying rhythm categorization

The automaticity of the neural categorization captured here could be suggestive of a low-level process rooted in nonlinearities of the earliest auditory processing stages. However, our results argue against this possibility. Indeed, the neural categories of rhythm identified here could not be explained by nonlinear transformations occurring at the earliest, peripheral stage of the ascending auditory pathway. Rather, our findings highlight a transformation from the representational geometry observed based on modeled auditory nerve responses, mainly tracking the physical temporal structure of the stimuli, toward the categorical representation measured with scalp EEG and consistent with behavioral responses.

Nonetheless, our data do not exclude that rudiments of this transformation could be found already in subcortical auditory nuclei, as has been proposed for the internal representation of periodic beat and meter elicited by rhythmic inputs structured according to an evenly spaced, isochronous grid of time intervals^61^. In fact, in line with perceptual representations in other domains^62^, this transformation could be hypothesized to emerge in the form of a gradient of sensitivity from physical to higher-level perceptual features across the processing neural network. For example, a cluster of high-level associative and sensorimotor cortices could support multimodal rhythm representations constrained by contextual and social semantics. These higher-level multimodal representations would thus guide the development and maintenance of categorical representations in sensory regions, for example through dynamic recursive exchanges of inputs^63^ (i.e., reentry^64^).

Our results are compatible with the view of a characteristic “warping” of the representational space where two rhythmic inputs are rendered more similar when they activate internal representation of the same category, as compared to physically equidistant rhythms internally assigned into different categories^65^. The results of the current study underscore the ability of the novel fRSA approach to locate boundaries delineating categories in this representational space, thus providing insights into their location and extent.

Here, and in line with a large body of prior behavioral work, we identified two rhythm categories: a smaller category encompassing rhythms including the 1:1 ratio, and a larger category, including the 2:1 ratio and spanning a broad range (between 1.3:1 and 2:1). Critically, the fact that we did not observe a boundary symmetrically splitting the representational space in two equal halves rules out the possibility that the two rhythm categories were driven by nonspecific range effects^66^.

A potential candidate to account for the observed rhythm categorization could be adaptation. While the absence of a categorical structure in the simulated auditory nerve responses argues against the role of fast low-level adaptation, a slower adaptation produced at later stages of the auditory pathway could, in principle, contribute to the neural categorization observed in scalp EEG activity^61,67,68^. Yet, the sharp dissimilarity in responses from each side of the category boundary as identified here would have to be driven by a highly nonlinear adaptation function that would abruptly begin to suppress the response to the tone starting the following rhythmic pattern when the corresponding inter-onset interval changes from roughly 290 to 260 ms (i.e., the shortest inter-onset interval duration found in the two conditions delimiting the category boundary). However, such adaptation mechanism has not been described yet in the literature^69–71^, and seems a priori too specific for a deterministic adaptation mechanism.

Instead, the observed categorical boundary might be compatible with a categorization process driven by detection of dissimilarity between the two intervals composing the rhythm^72^. At first glance, this may seem in agreement with previous behavioral research where two overarching categories of time intervals, even and uneven ones, have been consistently found to span the space of two-interval rhythms^14,19,41,73^. Accordingly, anisochrony detection, i.e., detection of deviance from evenness, might be one of the relevant mechanisms shaping the neural categorization of rhythm.

However, such a process of detection of deviance from isochrony would render the “uneven” category largely uniform over the possible two-interval rhythms, which is hardly compatible with the converging evidence for an underlying prototype or perceptual attractor located near 2:1 ratio (and possible other uneven attractors such as 4:3 and 3:1)^16–19,41,59^. Indeed, the neural responses recorded in the current study seemed to be systematically biased towards specific temporal templates also found in the behavior and in prior studies^16–19,21,59^. This result points towards a model where rhythm categories are encoded in prototypical patterns of neural activity^74^, rather than nonspecific adaptation or anisochrony detection.

All in all, delineating the specific representations underlying the rhythm categories identified in the current study will thus likely require going beyond electrical field potentials recorded with scalp EEG. For example, future work recording single neuron responses in the human temporal (as well as parietal and frontal) cortices appears a promising avenue to progress in our understanding of the neural processes supporting rhythm categorization^75–77^. The novel approach proposed here could constitute a major asset in this endeavor, by providing methodological tools to probe these neural responses across a diversity of signals.

### Rhythm prototypes

Categorical representational geometry has often been closely associated with the concept of a prototype^25,65^. In a Bayesian framework, prototypes can be thought of as modes in the prior distribution of rhythmic stimuli experienced over a lifetime^27,78^. Through probabilistic inference, these priors are then combined with information from sensory organs to support rhythm categorization^86^. In other words, the brain keeps track of what rhythms are encountered in the environment, and the representation of new rhythmic inputs is biased towards the rhythms that are more likely a priori^22,27^.

There is abundant behavioral evidence that perception of two-interval rhythms, such as the ones used in the current study, is pulled towards either prototypical 1:1 or 2:1 interval ratios^16–19,59^. An intrinsic attraction towards the mathematically simplest ratios is predicted by several models of rhythm perception, particularly by neurodynamic models of oscillatory networks, which propose that such biases emerge from predetermined physiological properties of the brain^26^.

However, previous tapping and perturbation detection studies in humans (see ^16^ for a review), as well as cross-species comparison research based on analysis of vocal communication recordings^49,50^, indicate that the 2:1 prototype may be, in fact, slightly shifted away from the smallest integer ratio. While most studies reported a shift in the direction of 1:1 ratio (i.e., “softening” of the contrast in duration between the two time intervals)^17,18^, others have found shift in the opposite direction (i.e., “sharpening” of the contrast in duration)^59^.

Our exploratory analysis of the similarity between continuous tapping responses and a set of prototypical rhythmic templates was broadly in line with a shift from exact integer ratios. Within the smaller category, the tapping dynamics mostly resembled a rhythm prototype with ratio 0.53, while the larger category featured tapping dynamics highly similar to both a softened (∼0.63) and a sharpened (∼0.68) prototype. It is worth noting that part of this effect may be explained by a pull towards the center of the range of rhythmic ratios presented in the condition continuum tested here. This possibility could be addressed in future work by adapting the rhythmic stimuli such as to locate the putative small-integer-ratio prototypes in the center rather than edges of the condition continuum.

Importantly, the tendency to converge towards a limited number of rhythm prototypes was also observed in the EEG responses. That is, there was a general alignment between the prototypes identified in tapping and in the neural responses, with observed prototypes compatible with bias away from mathematically simplest integer ratios. This result thus argues against a significant role of kinematic constraints in driving these more complex ratio prototypes. At the same time, EEG responses featured additional prototypes that were not encountered in the tapping responses. These prototypes included other complex ratios (0.59; approximately corresponding to 4:3), but also small integer ratios such as 0.50 (1:1) and 0.75 (3:1), the latter being in line with prior work probing the internal representation of rhythm indirectly via transient EEG responses elicited by expectation violations^79^.

Taken together, the heightened similarity to multiple rhythm prototypes per category observed in neural and behavioral data could reflect fingerprints of distinct underlying mechanisms (including individual differences^16^). The interplay between these mechanisms could further shape the rhythmic template eventually selected for perception and overt behavior during an explicit task. Some of these mechanisms might plausibly reflect a privileged role of mathematically simple rhythms emerging from generic oscillatory properties of neural assemblies^26,80^. Others could reflect plastic changes in the neural network acquired through experience (possibly via Hebbian learning^81^, and in line with recent Bayesian conceptualizations of rhythm perception^22,27,28^).

### Advantages of the approach and future directions

Building on recent advances in systems neuroscience^38^, a key advantage of the fRSA approach presented here lies in its capacity to capture categorization without taking potentially limiting or distorting assumptions about the exact form of the response that should reflect categorical information. In other words, the approach only relies on the fundamental definition of the categorization function^3–5^, which requires the system to generate selective (i.e., discriminant) responses to different categories, and reproducible (i.e., generalizable) responses when presented with the same category across a wide range of sensory conditions^1–5^.

As such, this approach goes a critical step beyond time-domain methods based on identification of time intervals between discrete events, which have proven difficult to apply beyond a small set of highly specific responses such as finger tapping^35,42^. Indeed, physiological responses (including brain activity), but also ecologically-valid movement elicited by rhythmic sounds (e.g., gestures or dance^82,83^) typically constitute complex trajectories including smooth fluctuations rather than series of transient events with clear temporal markers such as onsets ^37,82,84,85^. Therefore, we propose fRSA as a general method that could be applied to any continuous response elicited by periodic rhythmic stimulation, and that is, in principle, sensitive to any feature of the response that shows selectivity and reproducibility compatible with categorization.

Accordingly, the fRSA approach constitutes an important methodological advance, as it allows rhythm categorization to be probed directly from the dynamics of a wide range of neural responses (e.g. spiking rates, field potentials, oscillatory power fluctuations) at tempi that are ecologically valid (sometimes remarkably fast^86^), without relying on overt behavior, thus minimizing decisional, motivational, or motor confounds^16,87,88^. Moreover, as compared to other EEG approaches relying on transient responses elicited by expectation violations^79^, the fRSA approach allows to capture rhythm categorization (i) directly (i.e., without relying on additional processes and assumptions), (ii) by explicitly characterizing the representational geometry via dense sampling of the stimulus space (which is precluded by the long testing times typically required to assess EEG responses to deviant events), and (iii) without the need to estimate latency of the neural response^89^ (which is potentially ill posed especially at fast ecologically valid tempi).

This novel approach thus appears particularly well suited to address long-standing questions about the primitives and roots of musical rhythm, particularly the relative contribution of universal neurobiological constraints shared across species and culture-driven plasticity developing over the course of life through social learning. For example, it could allow to track how neural rhythm categories develop over the lifespan from birth, how they are shaped by cultural experience or body movement, and how this plasticity is supported by a network of brain regions shared in part by non-human species. Therefore, the framework developed in the current study appears promising to bridge the gap between recently found universality of some rhythmic structures in music on the one hand, and the vast inter-individual and cross-cultural diversity specific to human rhythm perception and production on the other hand.

## METHODS

### RESOURCE AVAILABILITY

#### Lead contact

Further information and requests for resources should be directed to and will be fulfilled by the lead contact: Francesca Barbero, Tomas Lenc, Sylvie Nozaradan.

#### Data and code availability

- Original data will be shared publicly upon publication.
- All original code will be shared publicly upon publication.

## EXPERIMENTAL MODEL AND STUDY PARTICIPANT DETAILS

### Participants

Eighteen participants (mean age ± SD = 26.0 ± 4.8 years, 13 females) were recruited in Brussels, Belgium. They reported various levels of musical and dance training (musical training, mean ± SD: 3.7 ± 6.2 years, range: 0-21 years, 11 participants never had any musical training; dance training, mean ± SD: 2.7 ± 3.9 years, range: 0-12 years, 9 participants never had any dance training). Sample size was estimated in accordance with previous studies using frequency-tagging to investigate rhythm perception^55,90–92^. All participants reported to have normal hearing and no history of neurological disorder. The experimental procedure was approved by the local ethical committee of the Saint-Luc Hospital - University of Louvain (project 2018-353) and conducted in accordance with the Declaration of Helsinki. Participants provided written informed consent to participate in the study (with 30 € of financial compensation).

## METHOD DETAILS

### Stimuli

The stimuli consisted of two-interval rhythmic patterns generated using Matlab R2022a (MathWorks). The two-interval rhythmic pattern was produced by presenting three auditory events (here: three identical tones) over time, while keeping the total duration of the pattern constant. In such a two-interval pattern, the first interval thus corresponds to the time between the onset of the first and the second event (*IOI1*, i.e., first inter-onset interval), while the second interval (*IOI2*) is defined as the time between the second and the third event. If the pattern is seamlessly looped, the third event of one pattern also constitutes the first event of the subsequent pattern (Fig. 1A).

The durations of the two intervals composing repeated two-interval patterns can be expressed as a ratio. For instance, if a given two-interval pattern exhibits a first interval that is twice as long as the second one, we can refer to that pattern as a 2:1 rhythm (Fig. 1A). In the current study, the position of the second tone was varied to obtain thirteen interval ratios equally spaced between 1:1 (which can also be expressed as the ratio between the first interval and the total duration of the pattern, i.e., 0.50) to 2:1 (i.e., 0.67). This thus yielded a 13-condition continuum, i.e., with a ratio granularity of thirteen increments (Fig. 1B, Table S1). We specifically focused on this 1:1-to-2:1 portion of the two-interval space, as this section has been shown to yield strongest categorical distortion in previous studies^22^, thus optimizing likelihood of capturing underlying neural correlates.

The thirteen two-interval rhythmic patterns were generated using 50-ms long pure tones with a carrier frequency of 300 Hz and a 10-ms linear onset/offset ramp. The patterns had a fixed total duration of 750 ms. This pattern duration was chosen since on the one hand it rendered unimanual tapping along with the stimuli reasonably comfortable for adults without musical training^47^, thus allowing comparisons of our results with the extensive previous work employing unimanual tapping tasks^16,18,19,21^. One the other hand, the chosen pattern duration was expected to yield stronger categorical distortions as opposed to slower tempi, as reported in previous behavioral studies^16^. Finally, for each of the thirteen conditions, the two-interval pattern was then seamlessly looped 30 times to form a 22.5 s-long stimulus sequence.

### Experimental procedure

The experiment consisted of 6 *listening blocks* and 3 *tapping blocks*, with the two types of blocks presented in alternation (a tapping block after every two listening blocks). In all blocks, the 13 different stimulus sequences were presented once in a randomized order.

During the listening blocks, participants were instructed to avoid any unnecessary movement and muscular tension, and fixate a cross displayed in front of them to minimize the presence of muscular and ocular artefacts in the EEG recording. Moreover, to ensure attention to the stimulus sequences, we used a task orthogonal to rhythm categorization whereby participants were required to detect transient volume drops in the sequences. The volume drops were obtained by decreasing the amplitude of 4 consecutive rhythmic patterns within a stimulus sequence to 85% of their amplitude. For each stimulus sequence, there could be either one volume drop (occurring in 2 out of the 6 presentations over all listening blocks), two volume drops (1/6) or none (3/6). After listening to the stimulus sequence without moving, participants verbally reported the number of detected volume drops and received immediate feedback.

During the tapping blocks, participants were instructed to tap in synchrony with the tones using the index finger of their preferred hand. Tapping was performed on a custom-made analog device (hereafter referred to as the ‘tapping box’) that was positioned by the participants’ side. Participants were instructed not to tap before the beginning of the stimulus sequences in order to obtain a valid period of baseline before trial onset (stimulus sequences were repeated when not meeting this criterion). Participants were also required not to wait too long to start tapping after the beginning of each stimulus sequence.

The experiment was implemented in Matlab R2016b (MathWorks, Natick, MA) using the Psychophysics Toolbox extensions^93–95^. The stimulus sequences were presented binaurally through insert earphones (ER-2, Etymotic Research, Elk Grove Village, IL; air-conducted sound from the level of the participant’s clavicle to decrease magnetic interferences) connected to a Fireface UC audio interface (RME Audio, Haimhausen, Germany; sampling frequency = 44100 Hz; 74.4 dB SPL). Participants performed the experiment while comfortably seated in a chair with their head resting on the chair support. The experiment had an overall duration of approximately 90 minutes including optional breaks between blocks.

### EEG recordings

We recorded brain activity using a 64-channels BioSemi Active Two EEG system (Biosemi, Amsterdam, Netherlands) with two additional channels placed on the left and right mastoids.

Recording sites included standard 10-20 system locations (channel coordinates can be found at https://www.biosemi.com/headcap.htm). Channel offset relative to the common mode sense (CMS) and driven leg (DRL) channel loop was kept below ± 50 mV (except in one participant in the main experiment whose offset was kept below ± 60 mV, but with no impact on the obtained signal-to-noise ratio of the expected responses).

An accelerometer was placed on the head of the participants to monitor whether participants complied with the instructions and avoided head movement during the listening blocks. The signals from all the channels and the accelerometer were digitized at a sample rate of 1024 Hz.

### Behavioral recordings

Tapping responses measured as tapping onsets and continuous force signal were recorded using the tapping box connected to the Biosemi Active Two EEG system’s Analog Input Box.

The surface of the tapping box was made of a conductive hard material, thus providing clear tactile feedback. While tapping also produced a small amount of auditory feedback, this was substantially attenuated by the ear inserts used to deliver the auditory stimuli (see below). The device recorded tapping onsets as moments in which the finger got in contact with the conductive surface and closed an electrical circuit. Simultaneously, the force exerted by the finger was recorded as a continuous signal using a six-axis force sensor (FT48224, ATI Industrial Automation, NC). The latency and jitter of the captured signals were below 1 ms, as measured with an oscilloscope.

The tapping onsets were digitized as triggers, while the force signal was digitized as the continuous signal coming from six different sensors of the tapping box at a sampling rate of 1024 Hz. Additionally, we also recorded a copy of the delivered acoustic signal through the Biosemi Active Two EEG system’s Analog Input Box to control for latency in the recording system, which was digitized at 1024 Hz.

### Auditory nerve modeling

To simulate responses elicited by the rhythmic stimuli in a set of auditory nerve fibers, we used an auditory nerve model developed by Bruce et al.^57^ as implemented in UR_EAR toolbox (version 2020b). Specifically, we modeled responses from 128 cochlear channels with characteristic frequencies logarithmically spaced between 130 and 3000 Hz. The default parameters used for cochlear tuning matched data available from human participants^96^. For each channel, we simulated a biologically plausible distribution of high, mid, and low-spontaneous-rate fibers (8, 12, and 31 respectively, i.e., 51 fibers in total)^97^. The resulting instantaneous firing rate was then summed across fibers and cochlear channels^98,99^, yielding an estimate of the variation in firing rate over the course of the 22.5-s long stimulus sequence in each condition.

## QUANTIFICATION AND STATISTICAL ANALYSIS

EEG and behavioral data were analyzed using Letswave 6, Letswave 7 (https://github.com/NOCIONS/letswave6, https://github.com/NOCIONS/letswave7) and custom-built scripts running on Matlab R2022a (MathWorks).

### EEG data preprocessing

A Butterworth highpass filter (4^th^ order, cut-off at 0.1 Hz) and lowpass filter (4^th^ order, cut-off at 64 Hz) was applied to raw continuous EEG data to remove slow drifts and responses at very high frequencies irrelevant to the current study. We subsequently downsampled the data to 256 Hz (i.e., by a factor of 4) to facilitate data handling and storage. We segmented the continuous data from -5 s to 27.5 s with respect to the onset of each stimulus sequence before performing artefact rejection. Following visual inspection of the data, we linearly interpolated noisy channels with the three closest neighboring channels (2 channels in 1 participant, 1 channel in 2 participants, no channels in the remaining 15 participants).

We then applied independent component analysis (ICA) to remove artefactual components due to blinks and eye movements. ICA matrices were computed from data preprocessed the same way as described above, except that we used a higher high-pass filter cut-off (1 Hz; 4^th^ order Butterworth filter) to improve artefact classification accuracy^100^. The data were re-segmented from 0 to 22.5 s with respect to the onset of the stimulus sequence (i.e., corresponding to each trial duration), and ICA matrices obtained using the probabilistic ICA model with Laplace approximation. Artefactual independent components (ICs) due to blinks and eye movements were identified from visual inspection of the ICs time course and topographies, and then removed (2 ICs removed in 3 participants, 1 IC removed in 14 participants, no IC removed in 1 participant). After this step, the preprocessing pipeline applied to the EEG data diverged depending on whether the data were then further analyzed in the frequency or in the time domain.

### EEG frequency domain analysis

After artefact rejection, the data were re-segmented from 0 to 22.5 s (i.e., total duration of individual stimulus sequences) relative to stimulus sequence onset. The duration of the resulting epochs thus corresponded to an exact integer multiple of the rhythmic pattern duration, hence preventing spectral leakage of responses at the frequencies of interest (determined as 1/pattern duration and harmonics) into the surrounding frequency bins after applying the Fourier transform^101^.

The data were re-referenced to average mastoids with the goal of maximizing the EEG responses to the acoustic stimuli^51,92^. The obtained epochs were then averaged in the time domain separately for each channel, condition and participant, to attenuate EEG activity not phase-locked with the stimulation. We then applied a Fast Fourier transform (FFT) yielding complex spectra for each channel, participant and condition with 0.044 Hz frequency resolution, corresponding to the difference in frequency between two consecutive frequency bins (i.e., 1/22.5 s stimulus sequence duration).

#### Feature extraction

In order to extract relevant features characterizing the neural response in each condition, we capitalized on the fact that the spectrum of any signal that is *systematically* repeated with a *fixed* repetition rate (i.e., periodically) will *only* contain peaks at specific frequencies corresponding the repetition rate (i.e., f = 1/repetition period) and its integer multiples (i.e., harmonics, 2f, 3f, 4f, etc.)^34,43,101,102^. In the current study, the stimulus sequences comprised of a seamlessly repeated rhythmic pattern presented with an exact fixed repetition period (i.e., 750 ms). Therefore, the spectrum of any response systematically elicited across repetitions of the rhythm pattern is expected to contain responses at frequencies corresponding to 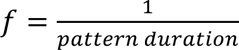 and harmonics (i.e., *f* = 1/0.75 s = 1.33 Hz, 2*f* = 2.66 Hz, …), which were thus considered as frequencies of interest in further analyses.

#### Significance of responses at frequencies of interest

Following a procedure adopted in previous frequency-tagging studies^44–46^, we first assessed the significance of the response at frequencies of interest at the group level. To do so, we computed the magnitude spectrum (i.e., absolute value of the complex spectrum) separately for each participant, condition and channel. We then computed the grand average across participants, conditions and all channels excluding the mastoids (i.e., 64 channels). Z-scores at the frequencies of interests were calculated as the difference between the magnitude at the frequency bin of interest and the averaged magnitude of 8 surrounding bins (4 on each side, excluding the immediately adjacent bins to avoid potential remaining spectral leakage), divided by the standard deviation of the same selected bins 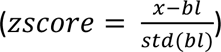. The response at a given frequency was considered significant if the corresponding z-score was higher than 3.09 (i.e., p < 0.001, one-tailed, signal > noise). Consecutive significant frequencies of interest were considered for the following analysis (see Fig. S3).

### EEG time domain analysis

After artefact rejection, we applied a fourth order Butterworth lowpass filter with a 10 Hz cut-off (i.e., to match the frequency range showing significant consecutive harmonics in the obtained EEG spectra, as measured using the z-score procedure described above and as further used in frequency-RSA analyses, see Fig. S3). The data were segmented from the onset to the end of the stimulus sequence (0-22.5 s) and re-referenced to average mastoids. The data were then further segmented to obtain successive chunks corresponding to the rhythmic pattern duration (i.e., 750 ms), and downsampled by a factor of 4 (i.e., to a 64-Hz sampling rate) to reduce dimensionality. This yielded a total of 180 epochs (6 trials x 30 pattern repetitions) per condition, channel and participant. To correct for possible offsets, the data was demeaned by subtracting at each time point within each epoch the average amplitude measured over the epoch for a given channel. The resulting epochs were then averaged separately for each condition, channel and participant.

### Behavioral data preprocessing

#### Inter-tap intervals

To analyze inter-tap intervals (ITIs), we adopted a procedure followed in previous sensorimotor synchronization studies^41,103^. First, we cleaned the series of tap onsets recorded as exact time stamps when the finger got in contact with the tapping box. The cleaning procedure was necessary since computing the ITI ratio for each repetition of the rhythmic pattern requires exactly three tap onsets. Accidental extra taps were removed by discarding tap onsets that were not separated from the preceding tap by at least 30 ms. Moreover, in order to discard responses that reflected attentional lapses or motor errors, we carried out the following steps. First, we corrected for the fact that humans tend to tap slightly before the pacing stimulus^47^. For each participant, we matched each tap onset with the closest tone and calculated the mean tap-tone asynchrony across all trials and conditions. We then subtracted the mean asynchrony from the onset time of each tap. All taps where the absolute value of the residual asynchrony to the closest tone onset was greater than 80 ms (i.e., less than half of the smallest inter-onset interval in the stimulus, which corresponds to the second interval in the 2:1 condition, 250 ms) were excluded from further analysis.

The obtained tap onsets were then used to calculate the ITIs separately for each repetition of the rhythmic pattern. Each tone in the given repetition was matched with the closest mean-asynchrony corrected tap. Then, we measured the time interval between the first and the second tap (*ITI1*) and the time interval between the second tap and the tap paired with the first tone of the directly following rhythm repetition (*ITI2*). The ITI ratio was calculated as 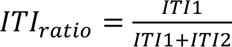. In case any of the tones was not matched with a tap, this precluded the computation of the ITI ratio and the whole rhythmic pattern repetition was excluded from further analysis. The computed ITI ratios were then averaged across pattern repetitions separately for each participant and condition.

#### Tapping onset time series

Separately for each participant, condition and tapping trial, we created a continuous time-domain signal with duration corresponding to the length of the stimulus sequence and 256 Hz sampling rate. The value of each sample corresponding to a tap onset time was set to 1 (i.e., a unit impulse) and 0 otherwise. Note that all tap onsets detected by the tapping box were used without any further preprocessing (i.e., unlike for the ITI analysis above).

#### Continuous tapping force

For each participant, condition and tapping trial, the continuous tapping force recorded from the six force sensors of the tapping box was segmented from -1 to 22.5 s with respect to onset of the stimulus sequences. For each sensor, the force signal recorded over the trial duration was baseline-corrected by subtracting at each time point the averaged signal over 1 s before trial onset to correct for potential offsets present in the recordings. The signal from the six sensors was then combined (using the device calibration matrix) to obtain the continuous tapping force orthogonal to the tapping box. The obtained tap-force signals were re-segmented from 0 to 22.5 s (i.e., stimulus sequence duration) relative to the onset of the stimulus sequences, and downsampled to 256 Hz.

### Behavioral data frequency domain analysis

The continuous responses (both continuous tap-onset time series and tapping force signals) were averaged across trials corresponding to different repetitions of the same condition, and an FFT was applied to obtain a response spectrum for each condition and participant. We assessed the significance of the responses at frequencies of interest at the group level (see Fig. S3). These computations were performed on the magnitude spectrum averaged across all participants and conditions, following the same steps as for the EEG responses.

### Behavioral data time domain analysis

#### Inter-tap intervals

The average ITI ratios were collapsed across participants and fitted either with a linear or a sigmoid model. Parameters were estimated by minimizing the least squares error, and the performance of each model was evaluated using leave-one-participant-out cross-validation. The sigmoid model was also fitted separately for each individual participant.

To test whether the tapped interval ratios overall significantly deviated from the stimulus ratios, we divided the range of stimulus ratios into 13 equal bins, computed histogram of the number of ITI ratios in each bin, and compared it to the one obtained under null hypothesis of a uniform distribution using the χ^2^ goodness of fit statistic.

#### Continuous tapping force

The continuous tapping force signals were also analyzed in the time domain, similarly to the EEG responses. To do so, a fourth order Butterworth lowpass filter with an 18 Hz cut-off was first applied to the preprocessed continuous tapping force signals (i.e., matching the frequency range whereby significant responses were identified using the procedure described above, see Fig. S3). Similarly to the EEG analysis, the data were then segmented into successive chunks of 750-ms duration, starting from onset time until the end of the stimulus sequence, thus yielding 90 epochs (3 trials x 30 pattern repetitions) per condition and participant, downsampled by a factor of 4, demeaned, and averaged separately for each condition and participant.

### Representational similarity analyses (RSA)

Neural and behavioral data were analyzed in the representational similarity analysis (RSA) framework^38,39^. RSA is a pattern information analysis comparing representational geometries that can be built from stimulus descriptors, empirical data, conceptual and computational models. Crucially, by moving from a unit-specific to a unit-free space, RSA allows to investigate whether a brain response reflects stimulus properties, higher-level categories or aspects of overt behavior. To characterize the representational geometries of the different types of data obtained in the current experiment, we computed several representational similarity matrices (RSMs) reflecting pairwise similarities across conditions. These RSMs were all characterized by a diagonal corresponding to the similarity between the signal and itself for each condition, thus with maximal similarity, separating the RSM into a lower and upper triangular half with strict symmetry between the halves.

#### Acoustic RSM

The acoustic RSM was obtained by taking the ratio between the first IOI and the total duration of the rhythmic pattern and computing the absolute difference of this value across all pairs of conditions. The resulting difference values were subtracted from 1 to yield a similarity matrix with ones on the diagonal. This RSM thus reflects the equal spacing of the rhythmic ratios along the condition continuum, i.e., a linear decrease of similarity across conditions (Fig. 1B).

#### Neural RSM

For each participant, a neural RSM was obtained as follows. First, the real and imaginary coefficients of the complex Fourier spectrum at the frequencies of interest (i.e., harmonics of the rhythmic pattern repetition rate where a significant response was observed at the group level) were extracted separately for each of the 64 channels. Then, separately for each condition, these values were concatenated into a feature vector, yielding number of dimensions equal to N frequencies of interest x 2 (real and imaginary coefficient) x 64 channels. Finally, we computed the similarity of these feature vectors using Pearson’s correlation across all pairs of conditions.

Additionally, we evaluated whether considering all 64 EEG channels and thus accounting for any individual differences in response topography is critical to reveal the categorical geometry of neural responses. To this end, we built the neural RSM from complex coefficients averaged across 9 fronto-central channels (F1, Fz, F2, FC1, FCz, FC2, C1, Cz, C2). Note that this is equivalent to extracting the coefficients from complex spectra of the channel-averaged time course. Channel selection was motivated by the observation that responses recorded over these channels seem to consistently capture EEG responses to repeating acoustic rhythms^36,54–56^ (see also Fig. S3).

Finally, a neural RSM was also built from time domain EEG responses. To do so, the preprocessed epochs corresponding to the average amplitude values measured over the time course of the rhythmic pattern were averaged across 9 fronto-central channels (F1, Fz, F2, FC1, FCz, FC2, C1, Cz, C2) separately for each participant and condition. Then, for each participant, the similarity of these time-domain responses across conditions was calculated using Pearson’s correlation across all pairs of conditions.

#### Behavioral RSM

Individual ITI RSMs were obtained by computing the absolute difference between the produced average inter-tap interval ratio across all pairs of conditions. To capture similarity, the difference values were subtracted from 1 as for the acoustic RSM.

Moreover, individual tap-onset RSMs were built, as for the neural responses, using feature vectors obtained by concatenating the real and imaginary coefficients at the frequencies of interest from the complex spectrum of the continuous tap-onset signals separately for each condition and participant (number of dimensions = 2 x N frequencies). The tap-onset RSM was then obtained for each participant by computing Pearson’s correlations between these feature vectors across all pairs of conditions.

Additionally, individual tap-force RSMs were obtained from continuous tapping force signal by following the same procedure as described for the tap-onset RSMs above.

Finally, as for the neural responses, individual tap-force RSMs were also built by comparing the time domain force values averaged across repetitions of the rhythmic pattern across all pairs of conditions using Pearson’s correlation.

#### Auditory nerve model RSM

As for the neural responses, the complex spectrum of the auditory-nerve model responses was calculated for each condition using FFT. In order to obtain an RSM directly comparable with the neural RSM, we extracted real and imaginary coefficients at the same frequencies of interest as for the EEG responses and concatenated them into a feature vector separately for each condition. The auditory nerve RSM was obtained by calculating Pearson’s correlation between feature vectors across all pairs of conditions.

### Theoretical models of rhythm categorization

Theoretical models of rhythm categorization were built based on the fundamental definition of a categorization function: maximal *similarity* of responses across conditions within the same category (thus setting pairwise similarity across these conditions to 1) and maximal *dissimilarity* of responses across different categories (thus setting the pairwise similarity of the corresponding conditions to 0).

Given that the stimuli employed in the current study are expected to elicit the perception of two rhythm categories^14,15,22,104^, we built a set of ten two-category models. The models differed from each other in terms of the position of the category boundary, i.e., in terms of the experimental condition where the switch from one category to the other would occur (Fig. 1E). Models where one of the categories spanned less than two conditions were excluded from the analysis, given that they cannot test within-category generalization. Rather than assuming the position of the category boundary a priori, having several theoretical models allowed us to estimate it from the data separately for each participant, thus accounting for any potential individual differences^16^. Likewise, we were able to assess whether the preferred theoretical models were consistent across participants and across neural and behavioral responses.

### Evaluation of shared structure in representational similarity matrices

#### Comparing response RSMs to categorical models

To investigate whether a response reflected rhythm categorization, we performed partial correlations between individual RSMs and the RSM of each theoretical model of rhythm categorization while partialling out the acoustic RSM to account for any representational structure driven by the stimulus (Fig. 1E). To this end, we calculated Spearman’s partial correlations between the lower triangular parts of the RSMs, excluding values from the diagonal to avoid inflating correlation values^105^. The best-fitting categorical model was defined as the one with the highest correlation coefficient.

Significance of the result for each individual participant was evaluated using permutation testing (5000 iterations), with the aim to probe whether the partial correlation coefficient with the best-fitting categorical model is higher than would be expected from chance. In each iteration, we randomly shuffled the values of the lower triangular response RSM (equivalent to shufling the condition labels) and computed the partial correlation between the shufled response RSM and the theoretical categorical models while partialling out the acoustic RSM. The correlation value corresponding to the winning theoretical categorical model identified from shufled response RSM was stored for each iteration. These values were used to build a null distribution of correlation values for statistical testing.

Significance was also assessed at the group level (permutation test, 10000 iterations) to test whether the group-averaged partial correlation coefficient with the participant-wise best-fitting model is significantly higher than expected from chance. For each iteration, we shufled all individual RSMs and found the best-fitting categorical model for each participant in the same way as described above. The average correlation coefficient of the best-fitting categorical model was stored to build a null distribution.

A p-value was calculated as the proportion of observations in the null distribution that had a correlation value higher than the one observed in the measured data^106^. To correct for multiple comparisons, p-values were adjusted using the Bonferroni correction for the 10 theoretical categorical models.

#### Comparing RSMs obtained from different measures

We were interested in comparing the RSMs obtained for different kinds of responses, such as ITI RSM, tap-onset RSM, tap-force RSM, and neural RSM. These comparisons were carried out using the following methods.

##### Comparing RSMs directly

To test whether two RSMs shared similar structure beyond what could be explained by the stimuli, we calculated Spearman’s partial correlation between the lower triangular parts of both RSMs while including the acoustic RSM as a covariate. Significance of the correlation was tested using a permutation test (5000 iteration), where partial correlation was computed from randomly shuffled RSMs on each iteration to build a null distribution of correlation coefficients. The same permutation procedure was employed to establish the significance of the average correlation coefficient across participants, i.e., a group level test (10000 iterations).

##### Comparing best categorical model fit

To test how prominent was the observed categorical structure at the group-level between two kinds of responses, the individual Spearman’s correlation coefficients obtained for the best-fitting model were Fisher-transformed, and further compared across the two kinds of responses using a paired t-test. In addition, Bayes factors were calculated quantify the evidence in favor of the alternative hypothesis over the null hypothesis (BF_10_), as implemented in the package bayesFactor for Matlab (https://github.com/klabhub/bayesFactor).

##### Comparing categorical boundary

In order to test whether two kinds of responses showed a similar category boundary position, we took into account the fact that theoretical models with similar category boundary positions were highly correlated. Hence, instead of relying on the best-fitting model, we considered all the possible models. For each response, individual Spearman’s partial correlation coefficients were obtained by correlating the corresponding RSM with each of the ten theoretical models of categorization. This yielded vectors of 10 correlation coefficients, which were further correlated (Spearman’s correlation) between the two kinds of responses separately for each participant. The obtained correlation values were then Fisher-transformed and tested against zero using a one-sample t-test, to test whether the correlation in the distribution of fit across category boundaries was similar between the two responses at the group level.

### Prototype analysis

In order to shed light on the structure of the categorical responses measured from brain and behavior, we evaluated the similarity of the continuous response (either tapping or EEG) in each condition and a set of prototypical signals. We built 76 time-domain prototypes with the same duration as the rhythmic sequences (i.e., 22.5 s). Each prototype was made of 30 seamlessly repeating 750-ms rhythmic patterns comprising two impulses arranged over time to create two inter-onset intervals (IOIs) with a given ratio. Across prototypes, the IOI ratios were equally spaced between a ratio of 0.50 (i.e., 1:1 ratio) to 0.84 (i.e., with contrast in duration between the two intervals sharper than the 3:1 ratio).

The similarity between each prototype signal and the response was evaluated at the group level using a bootstrapping procedure (1000 iterations). First, we obtained the magnitude spectrum of each prototype using FFT, and extracted a vector of magnitudes at the frequencies of interest, as selected for the analyzed responses (i.e., harmonics of the pattern repetition rate up to 8 Hz and 16 Hz for neural and tapping responses, respectively). Next, we used FFT to obtain the magnitude spectrum of the analyzed response separately for each condition and participant. To minimize the contribution of broadband noise to the magnitudes measured at the frequencies of interest, we applied a noise-correction procedure by subtracting from each frequency bin of interest the local noise baseline approximated as the average magnitude at 8 surrounding frequency bins (4 on each side, excluding the immediately adjacent bins to avoid potential remaining spectral leakage). Then, for each bootstrap iteration, we selected a random sample of 18 participants with replacement (i.e., at every iteration, each of the 18 participants could be selected multiple times). The vectors of magnitudes at the frequencies of interest were averaged across the sample of participants. For each condition, we correlated the average response magnitude vector with the vectors corresponding to each of the 76 prototypes, and stored the maximally correlated prototype, thus yielding a distribution of maximally correlated prototypes (Fig. 4A).

To test whether the distribution was concentrated near the stimulus ratio in each condition, we counted the maximally correlated prototypes inside a window centered on the stimulus ratio. The range of the window was defined by midpoints between successive ratios on the condition continuum. In other words, the window started half-way between the tested and the directly preceding condition, and ended half-way between the tested and the directly following condition. The counts inside the window were then compared to the counts outside of the window, in each case weighted according to the size of the respective range.

A similar approach was used to localize peaks in the distribution of maximally correlated prototypes. This was done by first splitting the conditions based on the category boundary identified using fRSA (median across participants) and building a marginal distribution separately for each subset of conditions. For each subset, local peaks in the pooled distribution were identified by sliding a window through the range of tested prototypes, with step size and width both corresponding to the spacing between neighboring stimuli on the condition continuum. The counts inside the window were compared to the counts falling in regions directly flanking the window (equivalent to 1/2 window width on each side).

Statistical significance was assessed using bootstrapping. The difference between weighted counts falling inside and outside the window of interest was calculated 1000 times from a resampled distribution of maximally correlated prototypes. P values were obtained based on the distribution of the bootstrapped data (Bonferroni corrected for multiple comparisons). Statistically significant peak inside the window was assumed if the difference values were greater than zero in more than 95% of the bootstrap iterations.

## Supplementary Information

**Table S1.** Inter-onset intervals (IOIs) used for stimulus construction. The table shows the interval (in seconds) between the onset of the first and the second tone (IOI 1), and between the second tone and the first tone of the following repetition of the rhythmic pattern (IOI 2). The table also shows the ratio between each IOI and the total duration of the rhythmic pattern. The values are shown separately for each of the 13 conditions, equally spaced between ratio 1:1 (equivalent to IOI 1 ratio 0.50) and 2:1 (equivalent to IOI 2 ratio 0.67).

| condition | IOI 1 (s) | IOI 2 (s) | IOI 1 ratio | IOI 2 ratio |
| --- | --- | --- | --- | --- |
| 1 (1:1) | 0.375 | 0.375 | 0.5 | 0.5 |
| 2 | 0.386 | 0.364 | 0.514 | 0.486 |
| 3 | 0.396 | 0.354 | 0.528 | 0.472 |
| 4 | 0.407 | 0.343 | 0.543 | 0.458 |
| 5 | 0.418 | 0.333 | 0.557 | 0.443 |
| 6 | 0.428 | 0.322 | 0.571 | 0.429 |
| 7 | 0.439 | 0.311 | 0.585 | 0.415 |
| 8 | 0.449 | 0.301 | 0.599 | 0.401 |
| 9 | 0.46 | 0.29 | 0.613 | 0.387 |
| 10 | 0.471 | 0.279 | 0.628 | 0.373 |
| 11 | 0.481 | 0.269 | 0.642 | 0.358 |
| 12 | 0.492 | 0.258 | 0.656 | 0.344 |
| 13 (2:1) | 0.503 | 0.248 | 0.67 | 0.33 |

**Table S2.** Summary of fRSA results separately for each indiidual participant. The results obtained with fRSA are shown separately for each participant and for the neural response (considering frequencies up to 8 Hz and 64 EEG channels) and for the tap-force response (considering frequencies up to 16 Hz). For each type of response, the table shows the (i) z-scored values of signal-to-noise ratio (zSNR), (ii) rhythmic ratio corresponding to the boundary of the best-fitting categorical model, (iii) Spearman’s correlation of the best-fitting categorical model RSM with the response RSM while partialling out the acoustic RSM, (iv) Bonferroni corrected p-value for the correlation obtained with a permutation test.

| participant | EEG |  |  |  | tap force |  |  |  |
| --- | --- | --- | --- | --- | --- | --- | --- | --- |
|  | zSNR | boundary | rho | P<sub>bonf</sub> | zSNR | boundary | rho | p<sub>bonf</sub> |
| 1 | 2.35 | 0.564 | 0.44 | 0.0079984 | 13.73 | 0.578 | 0.73 | 0.0019996 |
| 2 | 3.52 | 0.649 | 0.32 | 0.17796441 | 10.86 | 0.564 | 0.62 | 0.0019996 |
| 3 | 2.6 | 0.578 | 0.64 | 0.0019996 | 16.88 | 0.55 | 0.82 | 0.0019996 |
| 4 | 3.28 | 0.578 | 0.58 | 0.0019996 | 28.99 | 0.55 | 0.77 | 0.0019996 |
| 5 | 4.32 | 0.564 | 0.64 | 0.0019996 | 21.73 | 0.55 | 0.78 | 0.0019996 |
| 6 | 2.98 | 0.564 | 0.38 | 0.0379924 | 13.97 | 0.564 | 0.77 | 0.0019996 |
| 7 | 1.54 | 0.535 | 0.51 | 0.0019996 | 16.42 | 0.55 | 0.6 | 0.0019996 |
| 8 | 1.9 | 0.521 | 0.48 | 0.0039992 | 15.51 | 0.564 | 0.67 | 0.0019996 |
| 9 | 2.78 | 0.564 | 0.61 | 0.0019996 | 10.24 | 0.55 | 0.75 | 0.0019996 |
| 10 | 5.75 | 0.521 | 0.37 | 0.0439912 | 20.09 | 0.564 | 0.74 | 0.0019996 |
| 11 | 7.47 | 0.564 | 0.36 | 0.0619876 | 14.09 | 0.55 | 0.72 | 0.0019996 |
| 12 | 1.28 | 0.55 | 0.23 | 1 | 7.23 | 0.55 | 0.64 | 0.0019996 |
| 13 | 2.05 | 0.535 | 0.52 | 0.0019996 | 14.09 | 0.55 | 0.73 | 0.0019996 |
| 14 | 6.27 | 0.55 | 0.35 | 0.1139772 | 12.93 | 0.55 | 0.69 | 0.0019996 |
| 15 | 2.75 | 0.62 | 0.75 | 0.0019996 | 10.07 | 0.535 | 0.73 | 0.0019996 |
| 16 | 2.3 | 0.535 | 0.54 | 0.0019996 | 11.88 | 0.55 | 0.72 | 0.0019996 |
| 17 | 2.58 | 0.564 | 0.72 | 0.0019996 | 17.04 | 0.564 | 0.49 | 0.0019996 |
| 18 | 4.76 | 0.649 | 0.38 | 0.0359928 | 19.54 | 0.564 | 0.74 | 0.0019996 |

**Figure S1.**
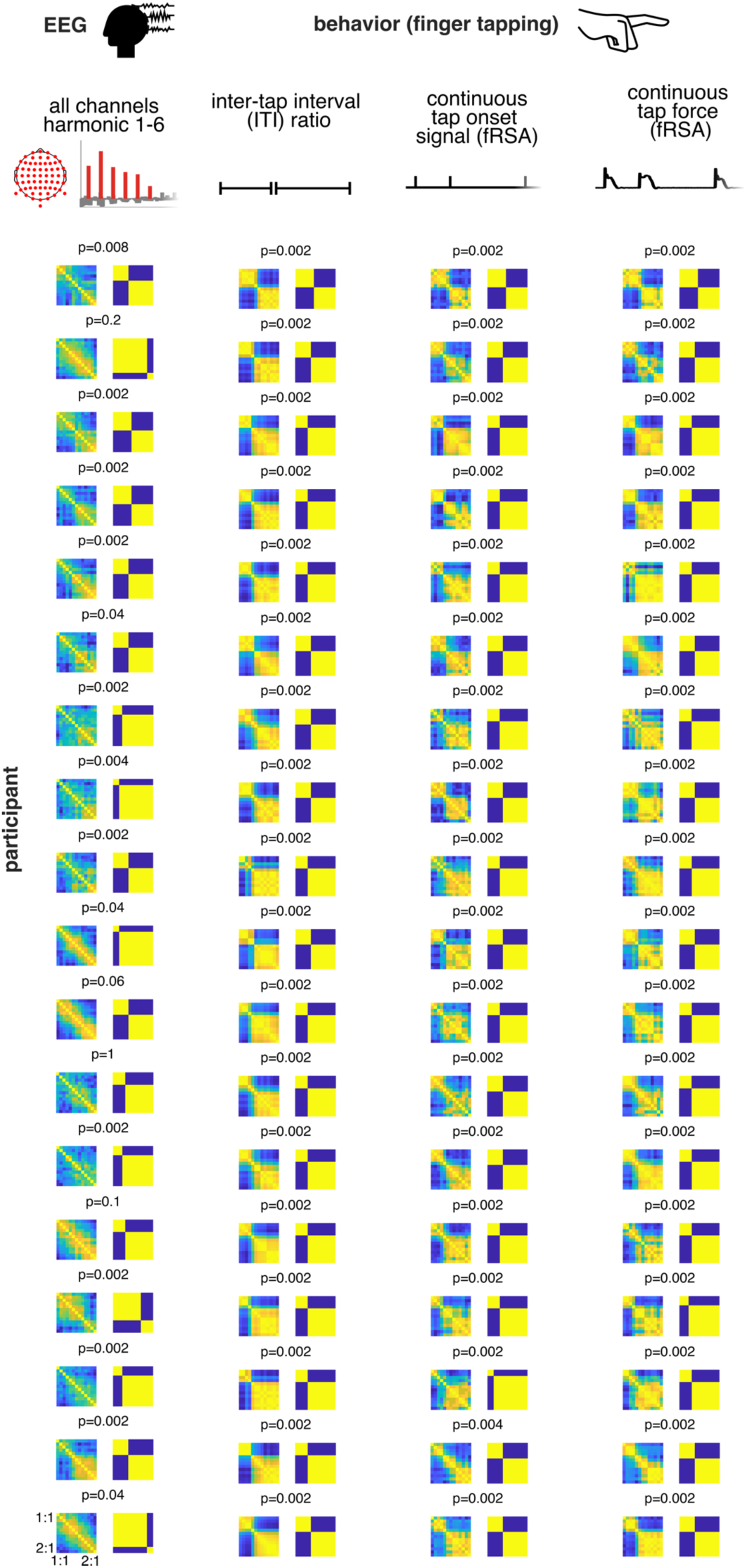
Representational Similarity Matrices (RSMs) of individual participants show overall consistent categorization and boundary locations. Each row corresponds to a single participant. Neural RSMs were obtained using fRSA by considering frequencies up to 8 Hz and all 64 EEG channels. The RSMs for ITI ratios were obtained from pairwise absolute differences in the produced ITI ratio across conditions. The RSMs for continuous tap onset and force signals were obtained with fRSA considering frequencies up to 16 Hz. Icon sources: “EEG” by Aenne Brielmann, and “finger” by iconcheese from the Noun Project under CC BY 3.0 license.

**Figure S2.**
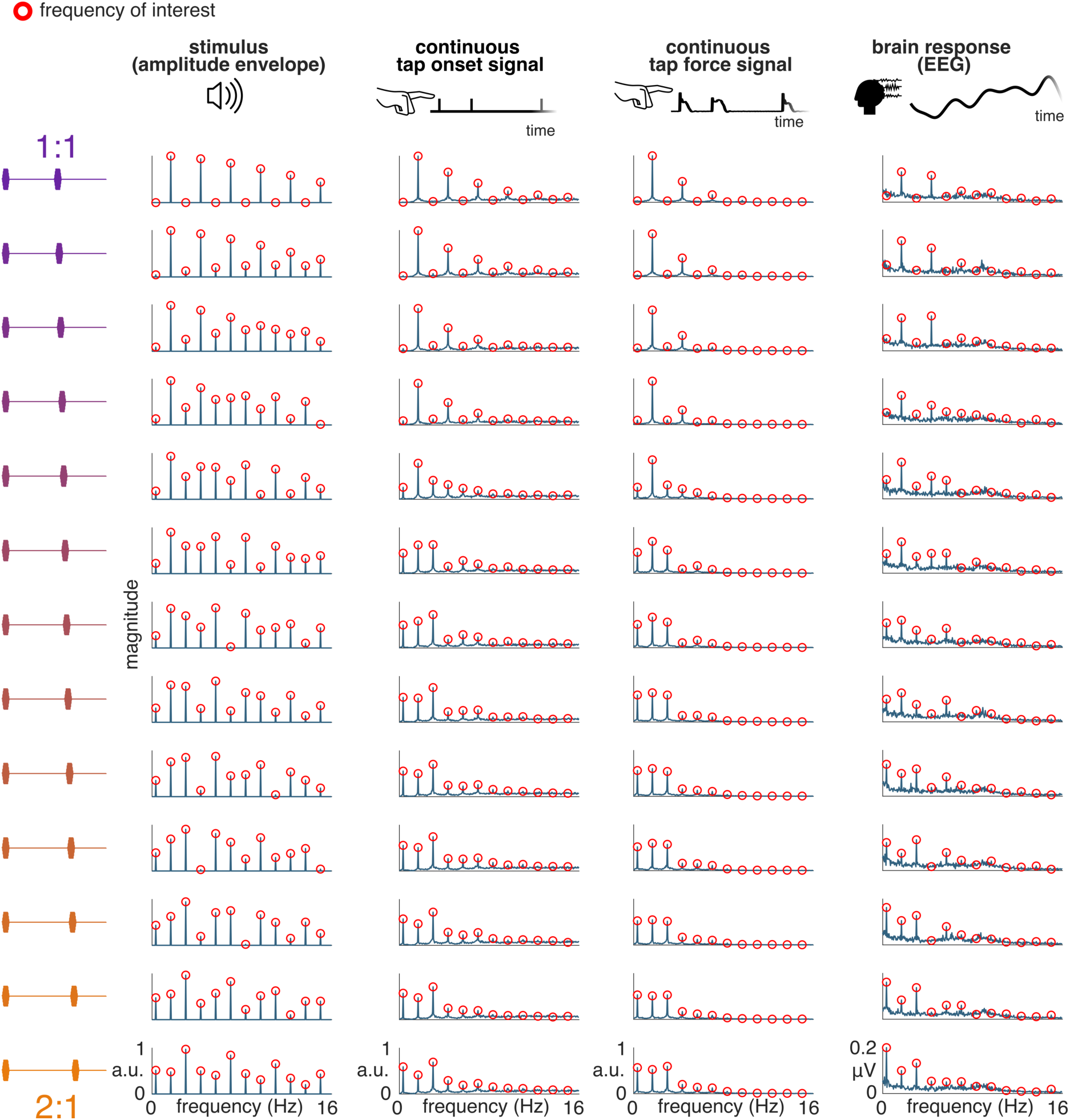
Grand average magnitude spectra of the stimulus, EEG, and tapping responses show peaks at a priori determined frequencies of interest. Grand average magnitude spectra shown separately for each condition. The stimulus spectrum is computed from the amplitude envelope of the auditory stimulus. Red circles highlight frequencies corresponding to the rate of rhythmic pattern repetition and harmonics. Icon sources: “sound” by PureSolution, “EEG” by Aenne Brielmann, and “finger” by iconcheese from the Noun Project under CC BY 3.0 license.

**Figure S3.**
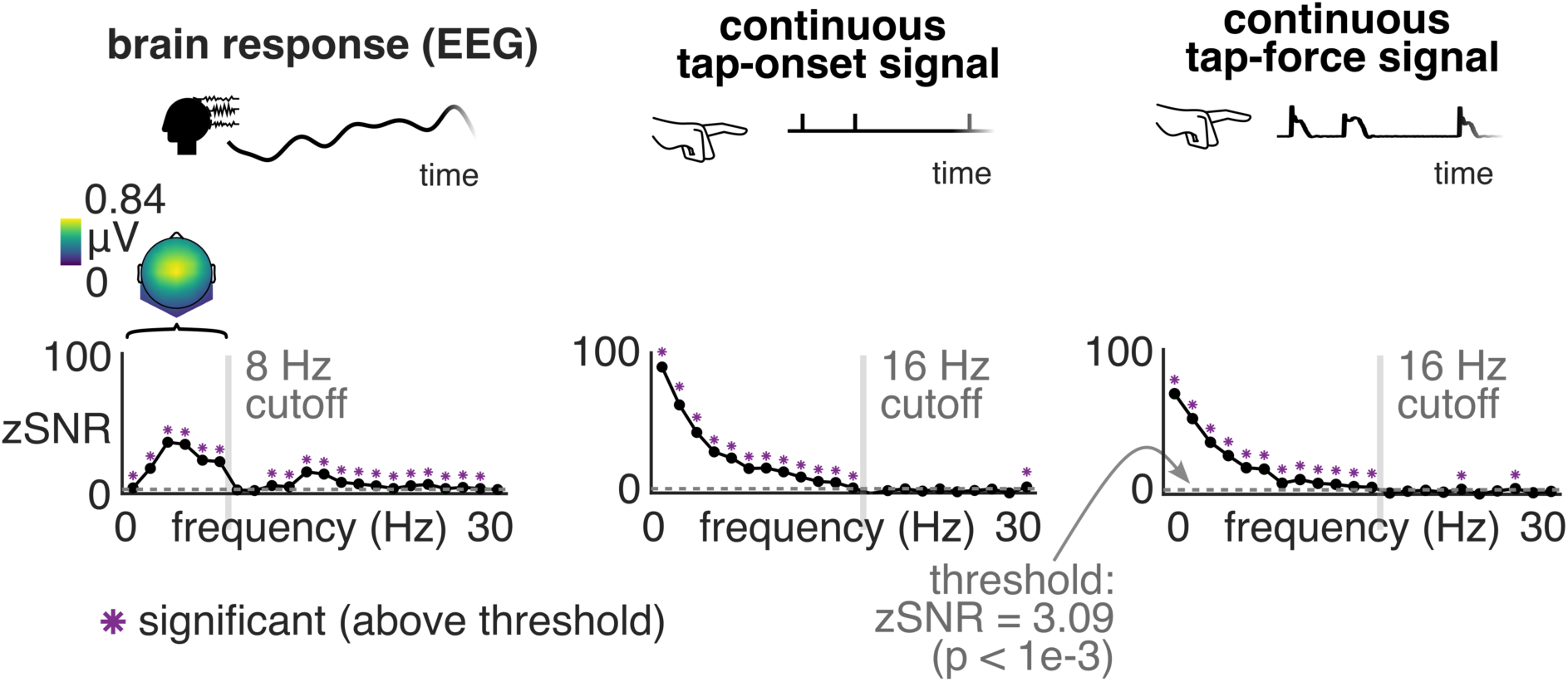
Significance of response harmonics. Z-scored values of signal-to-noise ratio (zSNR) of the response at each harmonic of the rhythmic pattern repetition rate obtained from the grand-average spectrum of the EEG and continuous tapping responses. The horizontal dashed gray line indicates the threshold corresponding to z-score 3.09 (equivalent to p < 0.001, one-tailed test, testing signal > noise). The zSNR value for each harmonic is shown as a black circle, and the magenta asterisk above indicates that the corresponding value is above threshold. A vertical grey line indicates the location of the cutoff used to build the RSMs (set to 8 Hz for neural and 16 Hz for continuous tapping responses). A topographical map of the magnitude summed across first 6 harmonics (i.e. up to 8 Hz, indicated by black brackets) is shown above the EEG plot. Icon sources: “EEG” by Aenne Brielmann, and “finger” by iconcheese from the Noun Project under CC BY 3.0 license.

**Figure S4.**
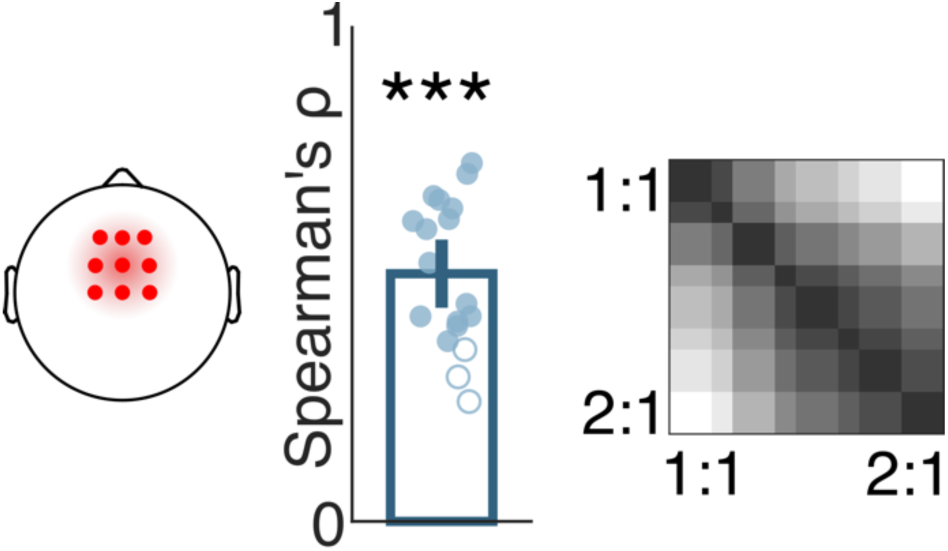
Similar categorization was obtained when applying fRSA to EEG responses averaged across fronto-central channels. Same as Figure 4, but with fRSA applied to a time course of the EEG response averaged across 9 fronto-central channels and considering frequencies up to 8 Hz. Blue circles show Spearman’s correlation of the EEG response RSMs with the best-fitting categorical model, obtained separately for each participant. Filled circles indicate a significant permutation test at the individual participant level, and asterisks indicate a significant permutation test at the group level (Bonferroni corrected, *p < 0.05, **p < 0.01, ***p < 0.001). Overlay of best-fitting significant categorical models across participants is shown on the right.

**Figure S5.**
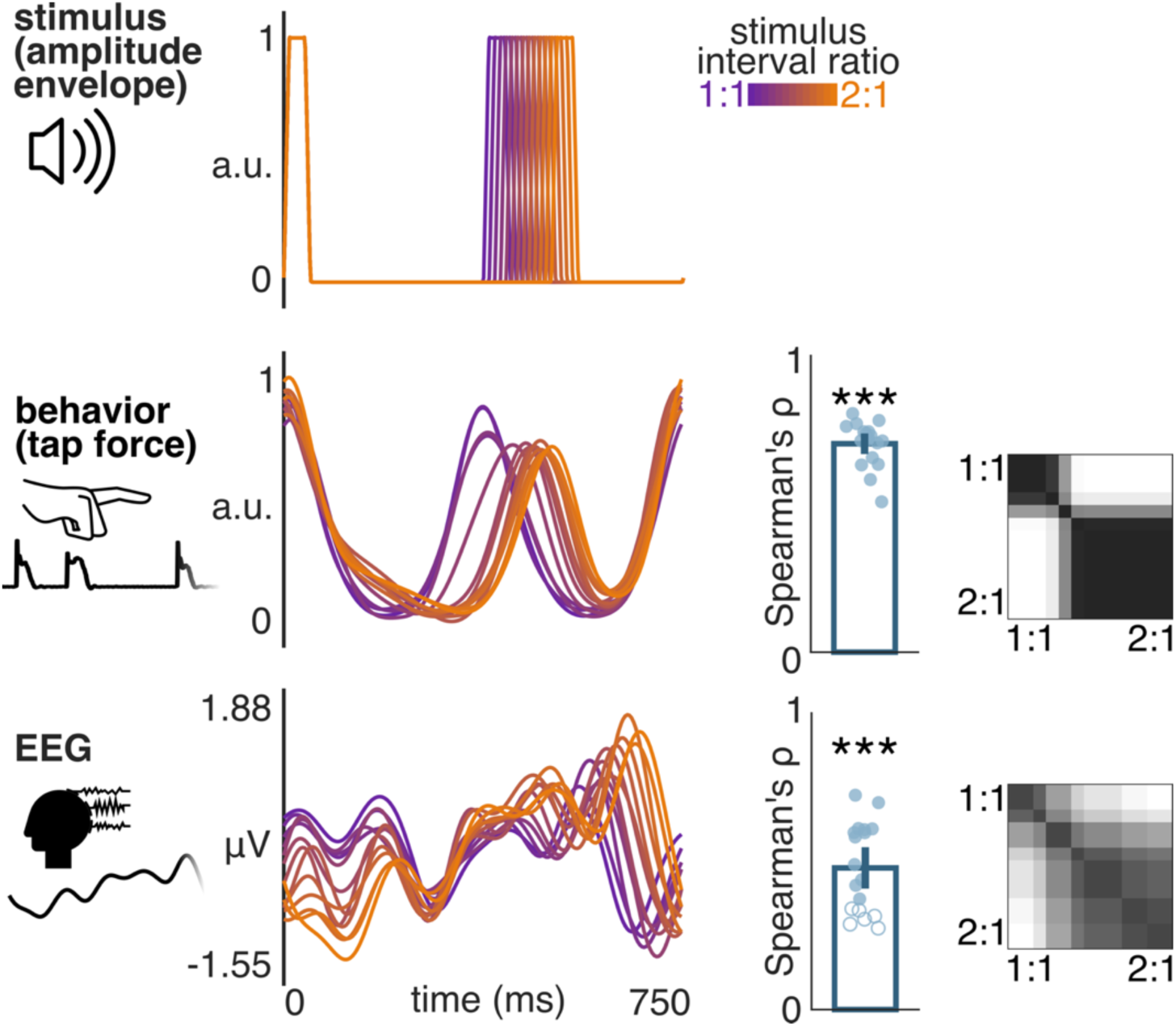
Similar categorization was obtained with time-domain analysis. The panel on the left shows an overlay of the grand-average time-domain response to a single repetition of the rhythmic pattern separately for each condition (indicated by the color gradient). The panel on the right shows (i) the correlation of a categorical model with the time-domain RSM based on the similarity of time-domain responses across conditions, and (ii) an overlay of best-fitting significant categorical models across participants. Filled circles indicate a significant permutation test at the individual participant level, and asterisks indicate a significant permutation test at the group level (Bonferroni corrected, *p < 0.05, **p < 0.01, ***p < 0.001). Icon sources: “sound” by PureSolution, “EEG” by Aenne Brielmann, and “finger” by iconcheese from the Noun Project under CC BY 3.0 license.

**Figure S6.**
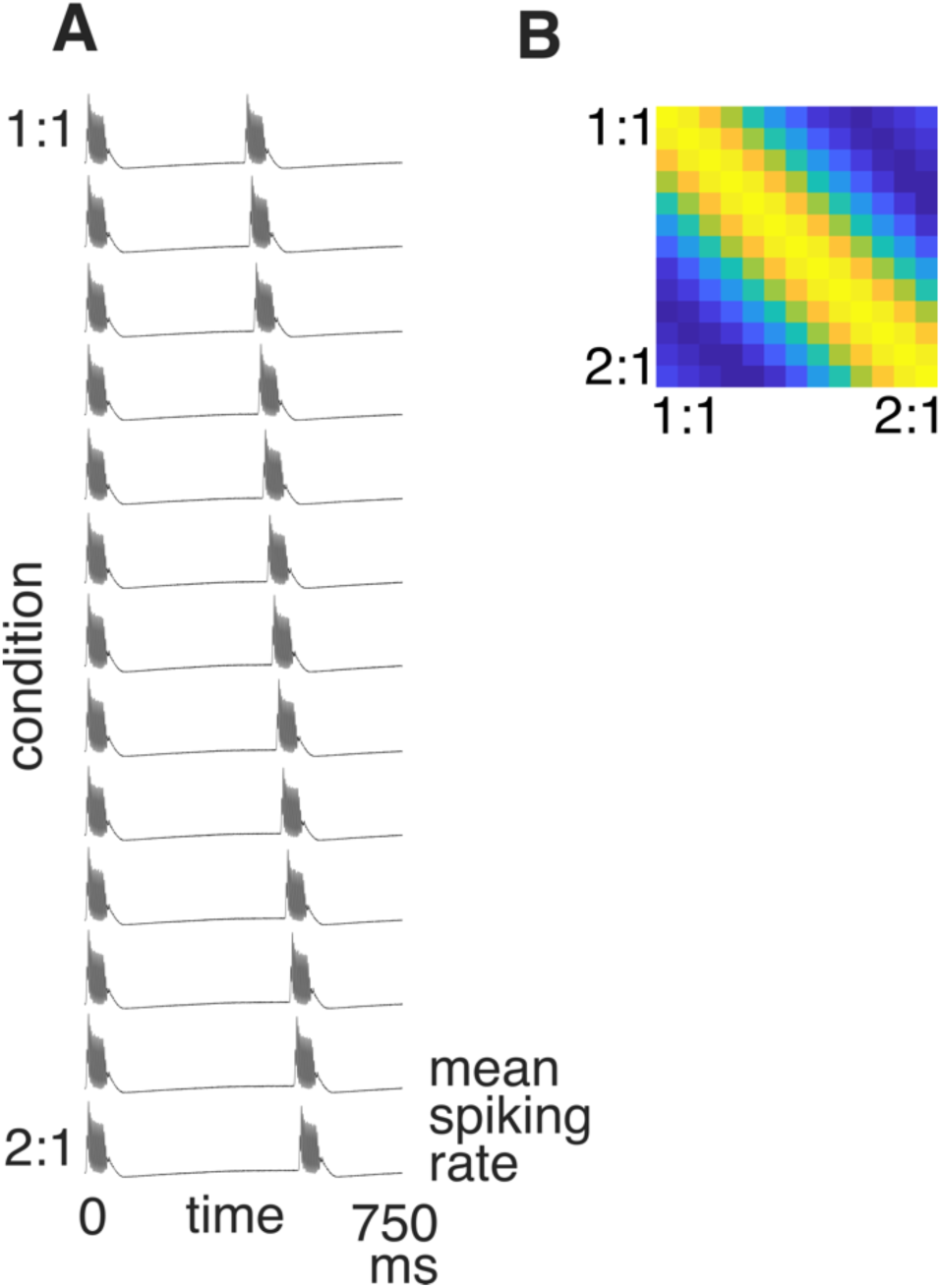
Auditory nerve model showing an absence of two-category geometry. A. Auditory nerve response to an average repetition of the 750-ms long rhythmic pattern separately for each condition. B. Auditory-nerve RSM built using fRSA and considering frequencies up to 8 Hz (i.e., same as for the EEG analysis).

